# A *SHR-IDD-PIN* regulatory network mediates minor vein differentiation in rice

**DOI:** 10.1101/2022.09.22.509058

**Authors:** Qiming Liu, Shouzhen Teng, Chen Deng, Suting Wu, Haoshu Li, Yanwei Wang, Jinxia Wu, Xuean Cui, Zhiguo Zhang, William Paul Quick, Thomas P. Brutnell, Xuehui Sun, Tiegang Lu

## Abstract

C_3_ and C_4_ grasses directly and indirectly provide the vast majority of calories to the human diet, yet our understanding of the molecular mechanisms driving photosynthetic productivity in grasses is largely unexplored. Here we define a genetic circuit comprised of SHR, IDD and PIN family members that specify vascular identify and ground cell proliferation in leaves of both C_3_ and C_4_ grasses. Ectopic expression and loss-of-function mutant studies of *SHORT ROOT* (*SHR*) paralogs in C_3_ *Oryza sativa* (rice) and C_4_ *Setaria viridis* (green millet) revealed a role in both minor vein formation and ground cell differentiation. Genetic and *in vitro* studies further suggest that SHR regulates this process through its interaction with Indeterminate Domain (IDD) IDD 12 and 13. We further show a direct interaction of these IDD proteins with a putative regulatory element within the auxin transporter *PIN5c* gene. Collectively, these studies indicated that a SHR-IDD regulatory circuit mediates auxin flow through the negative regulation of PIN protein expression to modulate minor vein patterning in the grasses.

## Introduction

Parallel leaf venation is a hallmark of grass leaves but the pattern of venation differs significantly between C_3_ and C_4_ grasses. A high vein density is a structural characteristic of nearly all C_4_ leaves (reviewed in Sedelnikova et al., 2018) and is necessary to support the metabolic cooperation of the two-cell photosynthetic system associated with C_4_ photosynthesis (Hatch and Slack, 1966). In both C_3_ and C_4_ grasses, it appears that the ontogeny of initial vein orders: midvein, major (lateral) veins and large intermediate veins (also known as rank-1 intermediate veins in C_4_) are largely similar (Ueno et al., 2006; Sedelnikova et al., 2018). However, as leaf expansion continues, C_4_ grasses initiate an additional file of small intermediate or small minor veins resulting in a greatly reduced interveinal distance (Ueno et al., 2006; Sedelnikova et al., 2018). Under hot, dry conditions C_4_ improvs photosynthetic efficiency largely due to a reduction in photorespiration and the need for less nitrogen and water to maintain high photosynthetic rates relative to C_3_ plants (Sage et al., 2012). In recent years, there has been much interest in engineering C_4_ traits into rice with the goal of doubling rice yields (Hibberd et al., 2008; Sage and Zhu, 2011). However, a key barrier to this program is a lack of fundamental understanding of the mechanisms that mediate a higher vein density in the C_4_ grasses.

Auxin has long been hypothesized to play a central role in the initiation and development of leaf veins (Sachs, 1969). Scarpella and colleagues showed that cell elongation is one of the first morphological markers of incipient vein formation and precedes a series of anticlinal cell divisions that are associated with the formation of provascular strands (Sawchuk et al., 2007). Expression of *PIN1* is one of the first molecular markers associated with the incipient vasculature in *A*. *thaliana* (Scarpella et al., 2006) and the grasses (Slewinski, 2013; O’Connor et al., 2014, 2017), suggesting that auxin flux (Sachs, 1981) together with auxin signaling (Verna et al., 2019) are essential regulators of vein initiation and development.

The *SHR/SCR* family has long been implicated in the development of C_4_ photosynthesis and mutations in this gene family disrupt the regular patterning of C_4_ leaves (Slewinski et al., 2013, 2014; Schlüter and Weber, 2020). The role of SHR is less clear in rice leaf tissues. Recent studies suggest that *OsSHR2* functions primarily in mediating stomatal differentiation in rice leaf tissue as overexpression (OX) of the maize ortholog of *OsSHR2* driven by a weak vasculature-specific promoter results in an increase in stomatal density (Schuler et al., 2018). However, strong constitutive expression of *OsSHR2* in rice results in dark green and curling leaf phenotypes suggesting a broader role regulating leaf development (Henry et al., 2017). Unfortunately, the nomenclature of the *SHR* family members is particularly problematic in the grasses. Initial reports of rice *SHR* genes focused on the gene expression profiles (Kamiya et al., 2003). However, subsequent studies from the Benfy lab revealed the *SHR* gene duplicated soon after the divergence of monocot and dicot lineages and suggested that the gene characterized as *OsSHR1* in the Kamiya et al. study is, in fact, *OsSHR2* based on sequence similarity to *AtSHR* in A.thaliana (Cui et al., 2007; Wu et al., 2014). Thus, it remains an open question what role if any OsSHR1 plays in regulating shoot or leaf development in rice. Supplementary Figure 1 and Supplementary Table 1 summarizes phylogenetic and functional relationships of the *Arabidopsis*, rice and maize *SHR* co-orthologs.

Here, we developed a series of transgenic lines overexpressing *SHR* orthologs to explore the function of SHR in grass leaf development. Phenotypic characterizations of multiple, independent single copy transgenic lines indicated *OsSHR1*, *S. viridis SHR1* (*SvSHR1*), *OsSHR2* and *SvSHR2* genes act redundantly to promote ground cell (e. g. mesophyll (M)) differentiation and repress bulliform cell development in rice and minor vein differentiation in both rice and *Setaria* tissues. Using CRISPR-Cas9, loss-of-function alleles *shr1* and *shr2* mutants in rice and *Setaria* alleles were generated and phenotypic characterizations of these plants were consistent with OX studies, as mutants had fewer mesophyll cell files between minor veins in both rice and *Setaria*. To identify additional components of the minor vein differentiation pathway, we generated loss-of-function mutants in *IDD* gene family members expressed in vascular tissues (Cui et al., 2013; Coelho et al., 2018) as IDD proteins have been shown to interact with the SHR/SCR complex (Hirano et al., 2017). Interactions of the IDD proteins with SHR using pull-down and BiFC assays revealed a direct interaction of IDD proteins with SHR. *osidd12* and *osidd13* double mutants resulted in fewer M cells between minor veins, consistent with a role of SHR-IDD in maintaining ground cell proliferation. Transcriptomic analysis of *osidd* mutants suggested that *PIN* family members were likely targets of IDD binding and here we provide *in vitro* and *in vivo* data showing a direct interaction of IDD 12 and 13 with a putative regulatory element within the rice *PIN5c* gene. Interestingly, in *OsPIN5c* OX lines, as few as three mesophyll cells could be found between adjacent minor veins and minor vein morphology. This finding suggests a role of *OsPIN5c* in promoting minor vein development and as a putative target to alter vein patterning in rice to more closely resemble that of C_4_ grasses. Finally, we propose a working model of minor vein differentiation driven by the SHR regulatory network in the grasses whereby OsPIN5c plays a critical role in the differentiation of both rank-1 and rank-2 intermediate minor veins.

## Results

### Ectopic expression of *SHR* mediates mesophyll cell proliferation and reduces vein density in the C_3_ plant rice and C_4_ plant *S. viridis*

To investigate the role of rice *SHR* orthologs in mediating vascular differentiation, we expressed the *OsSHR1* and *OsSHR2* cDNAs under the control of constitutive promoters resulting in T_0_ plants that seldom survived beyond the young seedling stage and produced few viable seeds (Supplementary Table 2 and Supplemental Figure 2). However, when native promoters were used to drive either *OsSHR1* (*OsSHR1::OsSHR1*) or *OsSHR2* (*OsSHR2::OsSHR2*), plants were qualitatively similar; slightly short-statured with adaxial leaf curling (Figure 1A, Supplementary Figure 3A - 3D, Supplementary Figure 4A, Supplementary Figure 5A - 5E and 5O). Histological characterizations of two independent single copy transgenic lines for *OsSHR1* and *OsSHR2* revealed a lower density of minor (rank-1 intermediate) veins, aberrant bulliform cell development and increased M cell number between minor veins relative to WT (Figure 1B, 1C, and 1D, Supplementary Figure 4B - 4F, Supplementary Figure 5F, 5G, 5H, 5Q and 5R). Interestingly, the density of major (lateral) veins (Supplementary Figure 4C, Supplementary Figure 5P) and the polarity and structure of vascular tissues (Supplementary Figure 3E - 3J and Supplementary Figure 5I - 5N) appeared to be unaffected by ectopic expression of *OsSHR1* and *OsSHR2*.

**Figure 1.**
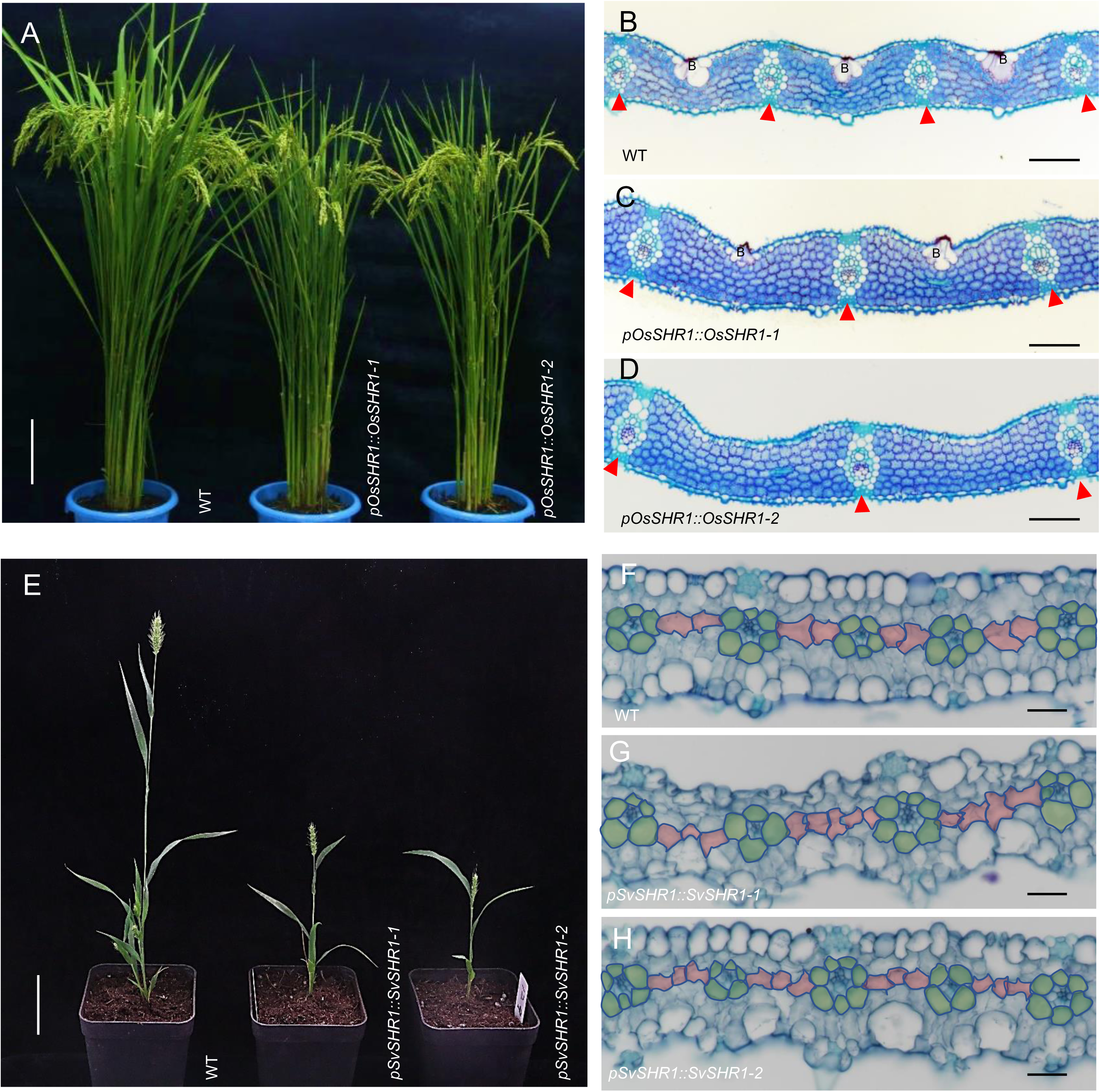
Ectopic expression of *SHR1* promotes mesophyll proliferation in C_3_ rice and C_4_ *Setaria viridis*. *OsSHR1* (LOC_Os07g39820) or *SvSHR1* (Sevir.2G383300) were overexpressed in rice cultivar Nipponbare (**A** - **D**) or *S. viridis* accession ME034V (**E** - **H**) driven by its native promoters. **A**, Mature plant phenotype of WT and two independent T_2_ *OsSHR1* OX lines at seed filling stage. **B** - **D**, Transverse paraffin sections in mid region of rice flag leaf (FL) showing increased mesophyll cells and smaller bulliform (B) cells in OX lines. WT shown in panel **B**, *pOsSHR1::OsSHR1-1* leaf shown in panel **C**, and *pOsSHR1::OsSHR1-2* (strong phenotype) leaf shown in panel **D**, red triangles indicate position of minor veins. **E**, Mature plant phenotype of WT and two independent T_2_ *SvSHR1* OX lines at seed filling stage. **F** - **H**, Transverse paraffin sections in mid region of *S. viridis* FL showing M cells (pink) and bundle sheath (green), WT shown in panel **F**, *pSvSHR1::SvSHR1-1* leaf shown in panel **G**, *pSvSHR1::SvSHR1-2* leaf shown in panel **H**. Bars for **A** = 20 cm, for **B**, **C** and **D** = 100μm, for **E** = 5 cm, for **F**, **G** and **H** = 100μm.

We also investigated the role of *S. viridis SHR* orthologs in mediating vascular differentiation in the leaf. Constructs overexpressing *SvSHR1* and *SvSHR2* cDNA driven by native promoters or constitutive promoters were introduced into *S. viridis* accession ME034V (Supplementary Table 2). Transgenic plants ectopically expressing SHR with native promoters were short-statured and set few seeds (Figure 1E and Supplementary Figure 6A). Cross sections and quantification of flag leaf cellular anatomy of two single copy transgenic lines showed a lower density of minor veins and increased M cell number between adjacent minor veins (Figure 1F, 1G and 1H, Supplementary figure 4G and 4I, Supplementary figure 6B, 6C, 6D, 6F and 6G). The density of lateral veins and the polarity and structure of vascular tissues were also unaffected through the ectopic expression of *SvSHR1* and *SvSHR2* (Figure 1F, 1G and 1H, Supplementary Figure 4H, Supplementary figure 6E). These results strongly suggest that *OsSHR1*, *OsSHR2*, *SvSHR1* and *SvSHR2* are functionally analogous and mediate mesophyll and minor vein density in both C_3_ and C_4_ grass leaf development.

Previous characterizations of ectopic *ZmSHR1* expression in rice, suggested a primary role for *OsSHR2* in mediating stomatal patterning (Schuler et al., 2018). To examine the role of OsSHR1 and OsSHR2 in epidermal cell fate decisions, we compared epidermal features in WT and OX lines through scanning electron microscopy (SEM). In WT plants there are typically two files of stomata between minor veins on the adaxial leaf surface. However, in transgenic lines, stomatal patterning is perturbed resulting in additional files of guard cells consistent with the increased interveinal distance, as well as occasional stomates occurring over the positions normally occupied by bulliform cells, which we never observed in WT plants (Supplementary Figure 7A, 7B and 7C). Stomatal complexes were also often reduced in size in OX plants (Supplementary Figure 7D, 7E and 7F). Interestingly, mesophyll lobing also appears to be reduced in OX lines (Supplementary Figure 7G, 7H and 7I). These traits suggest that SHR mediates either directly or indirectly the positioning of stomata in the epidermis as well as cell fate determination of mesophyll and bulliform cells (Figure 1B, 1C, and 1D, Supplementary Figure 4E and 4F, Supplementary Figure 5F, 5G and 5H).

### SHR mediates minor vein placement in rice and *S. viridis* leaves

To gain further insights into the role of endogenous SHR function, we constructed a series of loss-of-function alleles using CRISPR-Cas9 in both rice and *Setaria* (Supplementary Table 2 and Supplementary Figure 8). We generated loss-of-function alleles of *OsSHR1* and *OsSHR2* using gRNAs targeting the *OsSHR1* and/or *OsSHR2* coding region. T_2_ generation plants homozygous for *osshr1-1*, *osshr1-2*, *osshr2-1* and *osshr2-2* single mutants were morphological similar to WT. However, histological sections and quantification of flag leaf anatomy (Supplementary Figure 9) revealed that single *SHR* mutants reduce M cell differentiation and promotes minor vein initiation in the rice leaf, opposite to the *OsSHR* OX phenotypes and consistent with the hypothesis that OsSHR functions to promote M cell proliferation and inhibit vein initiation.

In rice, it was possible to generate *shr1 shr2* double mutants. Two T_2_ lines were identified, *osshr1-3 oshr2-3* were homozygous for a 353bp deletion in *OsSHR1*, and carried G and T insertions near the gRNA target site of *OsSHR2*. Line *osshr1-4 osshr2-4* was homozygous for a 352 bp deletion in *OsSHR1* and carried a single T insertion near the gRNA target site of *OsSHR2* (Supplementary Figure 8B). Translation from both alleles is predicted to occur at the same ATG codon as WT, but both *osshr1-3 osshr2-3* and *osshr1-4 oshr2-4* are predicted to encode frame-shifted and truncated proteins. Furthermore, we failed to detect transcripts in qPCR assays for either allele (data not shown). Together, these data suggest that *osshr1-3 oshr2-3* and *osshr1-4 oshr2-4* are complete loss-of-function alleles that are unlikely to encode any functional SHR1 and SHR2 product. Plants displayed a “loose” architecture with less compact tillers and a reduced stature relative to WT controls (Figure 2A) and leaves that curled in the opposite direction of *OsSHR* OX lines (abaxial leaf curling, Figure 2B and Figure 3A). The abaxial curling is consistent with enlarged bulliform cells and closer interveinal distances resulting in a higher density of bulliform cells on the adaxial surface (Figure 2C, 2D and 2E, Figure 3E and 3F). The average M cell numbers between minor veins varied across the leaf blade, with regions near the margin and midvein showing more variability. In WT plants, regions beyond one major vein distance away from the midvein and margins typically have 7 to 9 M cells between minor veins, whereas minor veins in corresponding regions of *osshr1 oshr2* lines were typically separated by 3 to 5 M cells (Figure 2C, 2D, 2E and Figure 3B). In contrast, the total number of minor veins increased in *osshr1 osshr2* relative to WT (Figure 3D), In addition, BS cell files were often disrupted with intervening M cells, and sclerenchyma cells that are typically positioned at the adaxial and abaxial surface of veins, were often replaced by a file of M cells (see arrows in Figure 2D and 2E).

**Figure 2.**
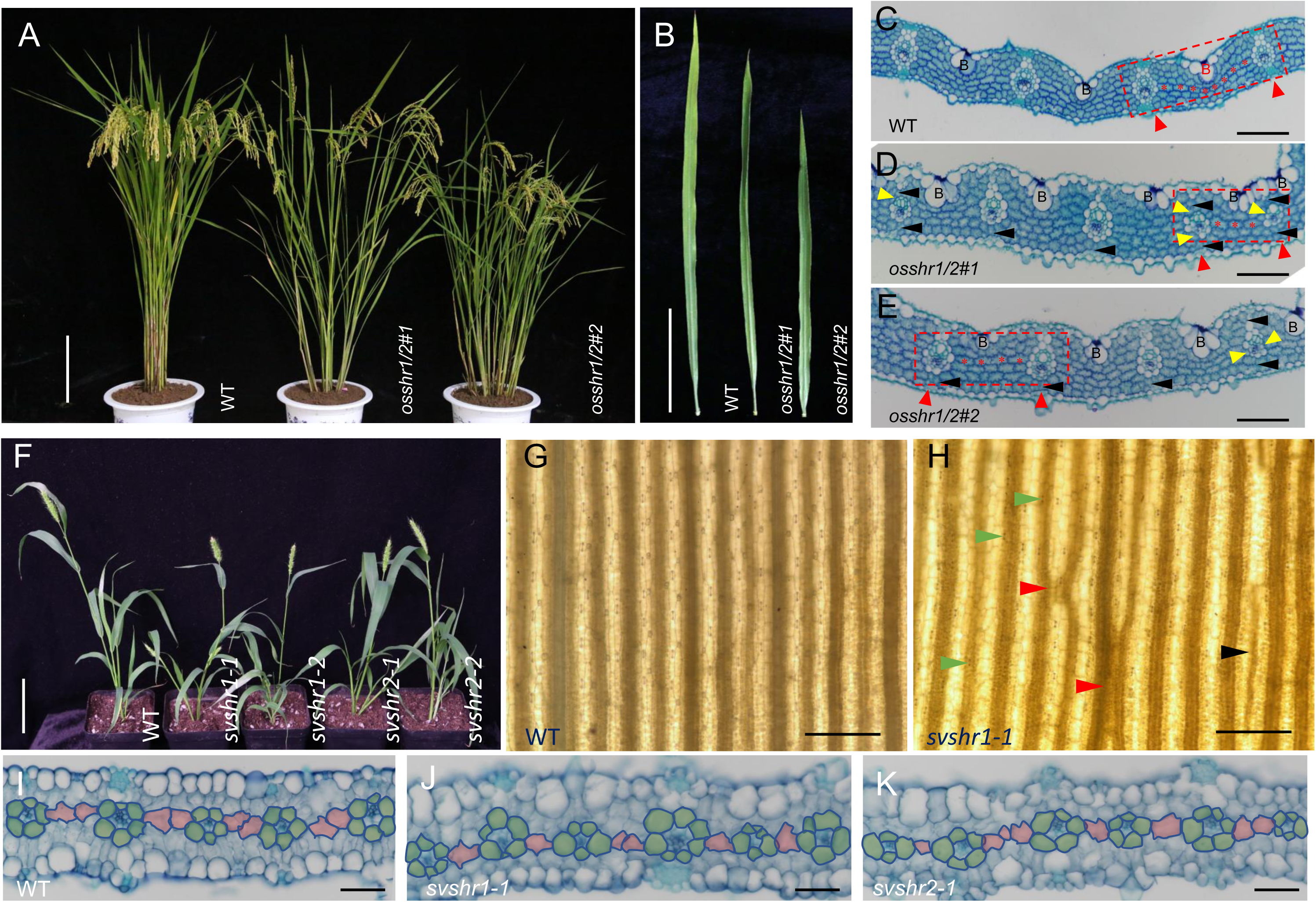
Loss of function SHR alleles condition altered vein patterning in rice and *Setaria*. *OsSHR1*, *OsSHR2* (LOC_Os03g31880), *SvSHR1* and *SvSHR2* (Sevir.9G361300) loss-of-function double (*osshr1*/*2*#*1* presents *osshr1*-*3 osshr2*-*3* and *osshr1*/*2*#*2* presents *osshr1*-*4 osshr2*-*4*) or single (*svshr1-1*, *svshr1-2*, *svshr2-1* and *svshr2-2*) mutants were generated by CRISPR-Cas9 in Nipponbare or ME034V, respectively. **A** and **B**, Mature plant phenotype of WT and two independent *osshr1 osshr2* double mutants at seed filling stage (**A**) and abaxial leaf surface of FL (**B**). **C**, **D** and **E**, Transverse paraffin sections of FL showing M cells, B cells and minor veins, red triangles indicate the minor vein position and red rectangles highlight the region between two adjacent minor veins with M cells indicated by red asterisks, black solid arrows show missing sclerenchyma cells and yellow solid triangle indicate a M cell within a file of BS cells (**D** and **E**). **F**, Independent *S. viridis svshr1* and *svshr2* single mutants were shorter than WT plants at seed filling stages in T_2_ generation plants. **G** and **H**, Light micrographs of cleared ME034V leaf tissue (I-KI stained to reveal starch) showing that the regular parallel vein pattern (WT, **G**) is disrupted in mutant leaves (**H**), black solid triangle shows incomplete Kranz anatomy along a minor vein, red solid triangle shows two adjacent minor veins without intervening M cells, and green solid triangles indicating ectopically located stomata complexes. **I**, **J** and **K**, Transverse paraffin sections in mid region of ME034V FL showing M cells (pink) and bundle sheath (green) in WT (**I**), *svshr1-1* (**J**) and *svshr2-1* (**K**). Bars for **A** = 5 cm, for **B** = 20 cm, for **C**, **D** and **E** = 100μm, for **F** = 5 cm for **G** and **H** = 1 mm, for **I**, **J** and **K** = 100 μm.

**Figure 3.**
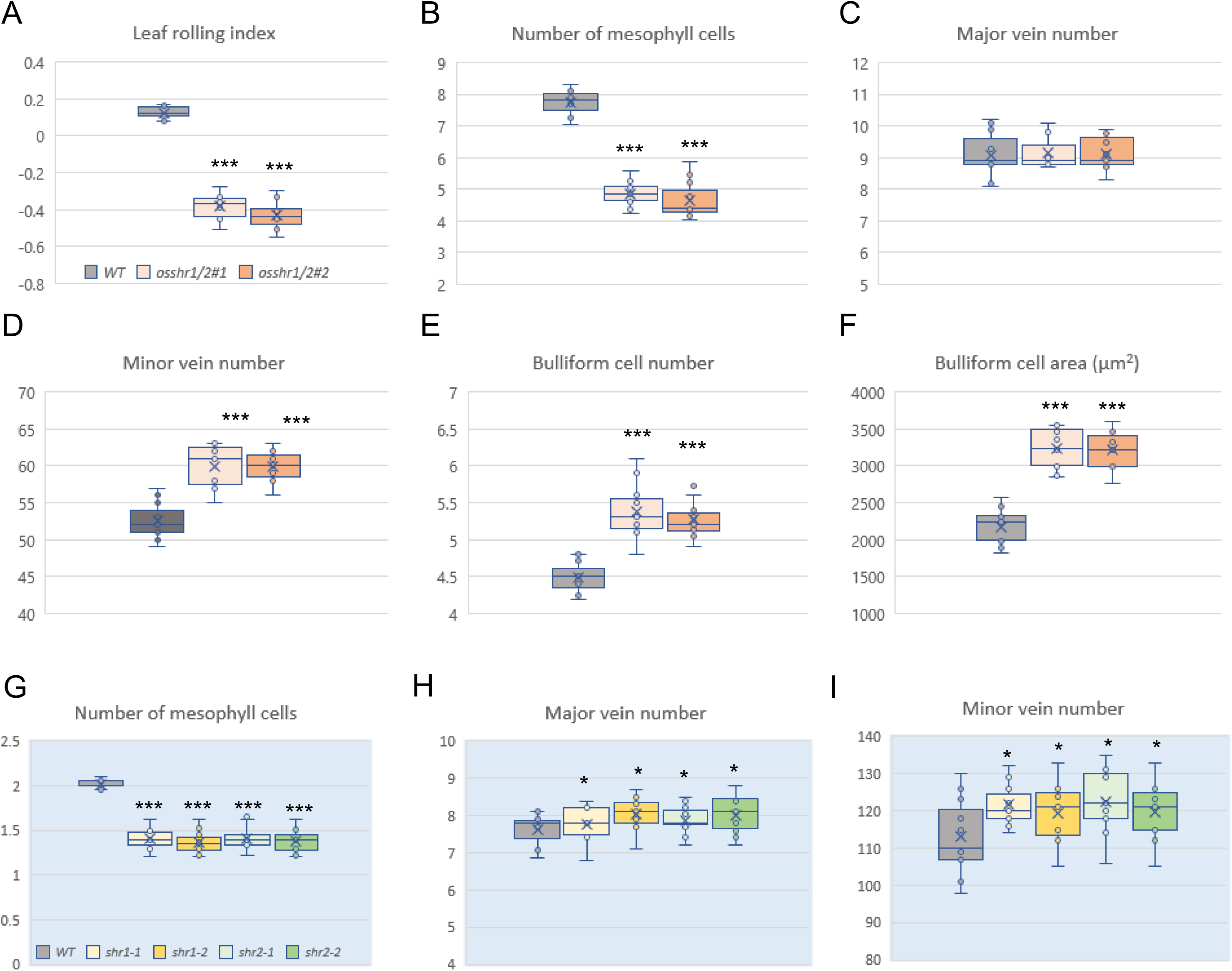
Quantification of leaf traits in *SHR* mutants of rice and *S. viridis*. Leaf trait data in rice WT and independent *osshr1 osshr2* double mutants (**A - F**) and in ***S. viridis*** WT and independent *shr* single mutants (**G, H** and **I**). P values calculated using student’s t-test, * p<0.05, *** p<0.001, n = 11 for each genotype.

We also generated loss-of-function alleles of *SvSHR1* and *SvSHR2* using gRNAs targeting the *SvSHR1* and/or *SvSHR2* coding region to generate alleles (Supplementary Figure 8C). Plants homozygous for *svshr1-1*, *svshr1-2*, *svshr2-1* and *svshr2-2* are morphologically similar with a reduced stature and poor seed set relative to WT (Figure 2F). Light micrographs of cleared leaf tissue (I-KI stained to reveal starch) revealed a disruption in the typical parallel vein pattern that included incomplete Kranz anatomy, adjacent minor veins, and ectopically located stomata cells (Figure 2G and 2H). Histologic sections (Figure 2I, 2J and 2K) and quantification of M cell numbers between minor veins revealed an increase in minor vein number (Figure 3G, 3H and 3I). We were unable to recover seed from double mutant *svshr1svshr2* individuals. Collectively, these data indicate that in addition to controlling M cell proliferation and vein patterning, SHR1 and SHR2 regulate both BS and sclerenchyma cell identity in grasses.

### *OsSHR1* and *OsSHR2* expression is enriched in vasculature

Quantitative PCR characterizations of *OsSHR1* and *OsSHR2* expression indicated that both genes are expressed in vegetative organs including root and shoot tissues (Figure 4A). *In situ* analysis of developing leaf tissues revealed expression of *OsSHR1* in vascular files of P2 - P5 leaf primordia and relatively weak expression in the subepidermal developing sclerenchyma cells adjacent to all vein ranks (see white arrow in Figure 4B). However, signal was less intense in P5 leaf primordia and not detectable in P6 blade tissues (data not shown). Signal was most apparent in the midvein of P2 leaves, major veins of P3 - P5 leaves and in some minor vein initials in P3 - P5 leaf primordia (Figure 4B), in a pattern that is qualitatively similar to *OsSHR2* expression (Schuler et al., 2018). To further define the expression domains of rice *SHR*, we generated a promoter::GUS reporter construct with the identical *OsSHR1* and *OsSHR2* promoter fragments as used in the OX studies (Supplementary Table 2). Lines expressing the GUS protein displayed strong vascular expression in shoot and root primordia and root and leaf tissues, with expression in root tissues highest in root tip and incipient lateral root initials (Figure 4D - 4M).

**Figure 4.**
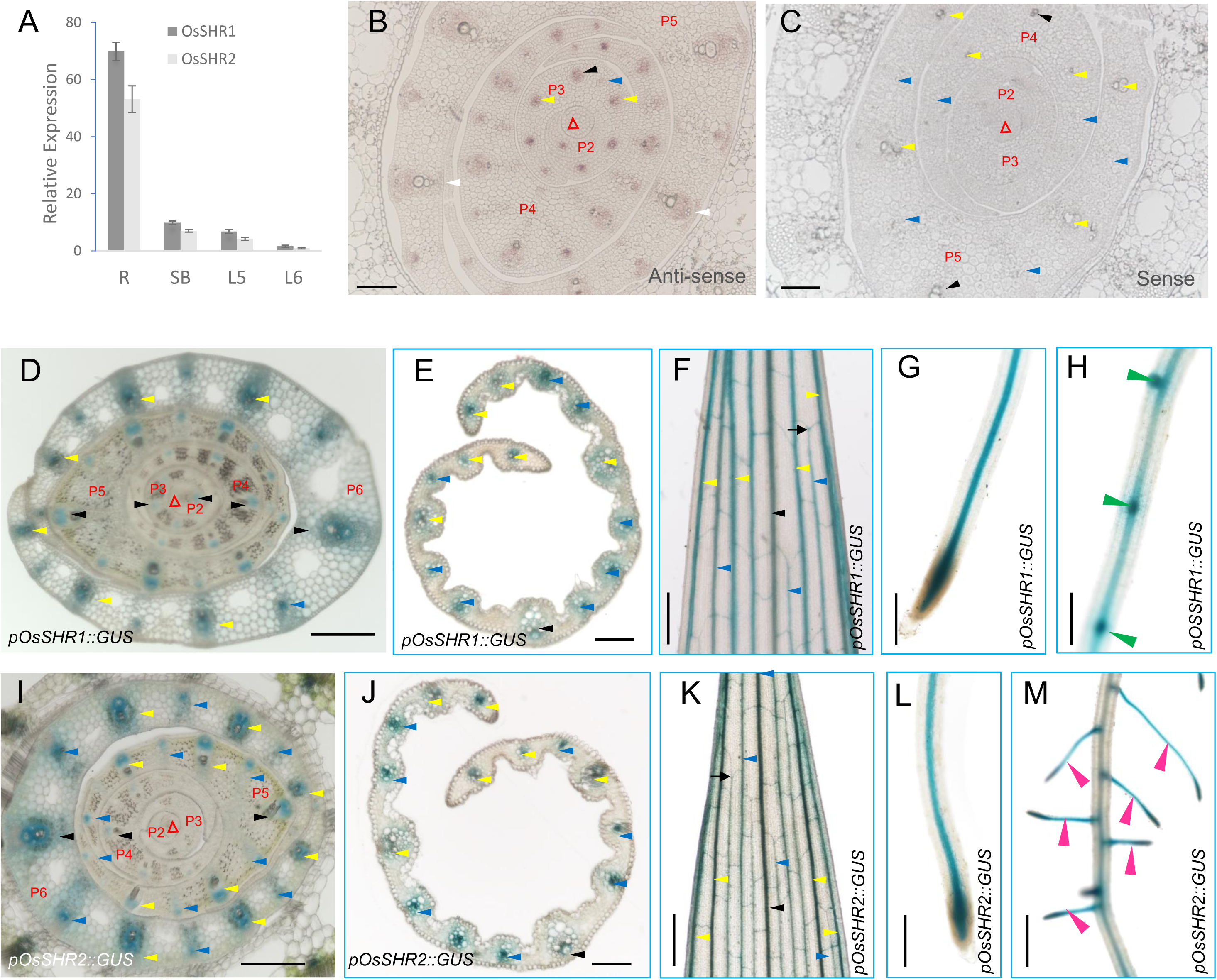
*OsSHR1* and *OsSHR2* expression is enriched in vasculature. **A**, Quantitative reverse transcription PCR analysis showing *OsSHR1* and *OsSHR2* expression in the root (R), shoot base (SB), the fifth leaf (L5, fully expanded) and the sixth leaf (L6, new emerging) in 20 DAG seedlings. **B** and **C**, *in situ* analysis of *OsSHR1* showing expression in developing vasculature of P2 to P5 leaf primordia as detected by anti-sense *OsSHR1* probe (**B**) and sense probe as control (**C**). **D** - **H**, Gus-staining of *pOsSHR1::GUS* transgenic plants including cross section of shoot base (**D**), cross sections of L5 leaf tip (**E**), L5 leaf tip (**F**), root tip (**G**) and lateral root initiation sites (**H**) of 20 DAG seedlings. **I** - **M**, Gus-staining of *pOsSHR2::GUS* transgenic plants including cross section of shoot base (**I**), cross sections of L5 leaf tip (**J**), L5 leaf tip (**K**), root tip (**L**) and lateral roots (**M**) of 20 DAG seedlings. White arrows indicate subepidermal developing sclerenchyma cells adjacent to veins **(B**), red triangles indicate the SAM in panel **B** - **D** and **I**, black arrows Indicate the developing midribs, yellow arrows indicate the developing major veins and blue arrows indicate the developing minor veins in panel **B** to **M**, green arrows indicate the initiating lateral roots (**H**), and pink arrows indicate developing lateral roots (**M**). Bars for **B** and **C** = 50 μm, for **D** and **I** = 200 μm, for **E** and **J**= 1 mm, for **F** and **K** = 0.5 mm, for **G**, **H** and **L** = 2 mm, for **M** = 1 mm.

### OsSHR1 directly interacts with OsIDD12 and OsIDD13

Structural characterizations of the SHR-SCR heterodimer revealed a direct interaction with an IDD domain protein (Hirano et al., 2017). Thus, we examined the expression and function of several IDD gene family members predicted to be expressed in vascular tissues in *Arabidopsis* and rice (Cui et al., 2013; Coelho et al., 2018) and identified *OsIDD12* and *OsIDD13* as vasculature specific (Figure 5). Promoter::GUS reporters were created with both OsIDD12 and OsIDD13 (Supplementary Table 2) and GUS-stained leaf and primordia tissues indicated that *OsIDD12* (Figure 5A and 5B) and *OsIDD13* (Figure 5C and 5D) expression was largely restricted to vascular tissue of the expanding leaf tips and in P5 and P6 leaf primordia at the shoot base. Further *in situ* analysis using anti-sense probes to *OsIDD12* (Figure 5E and 5F, Supplementary Figure 10A) and *OsIDD13* (Figure 5G and 5H, Supplementary Figure 10B) indicated that *IDD* genes were expressed in developing veins of leaf primordia. Collectively, our data revealed *OsIDD12* and *OsIDD13* are expressed in the same tissues and with similar developmental timing as *SHR* paralogs in vascular tissues of developing leaves. Thus, rice *IDD12* and *IDD13* represent likely targets of SHR interaction in leaf tissue.

**Figure 5.**
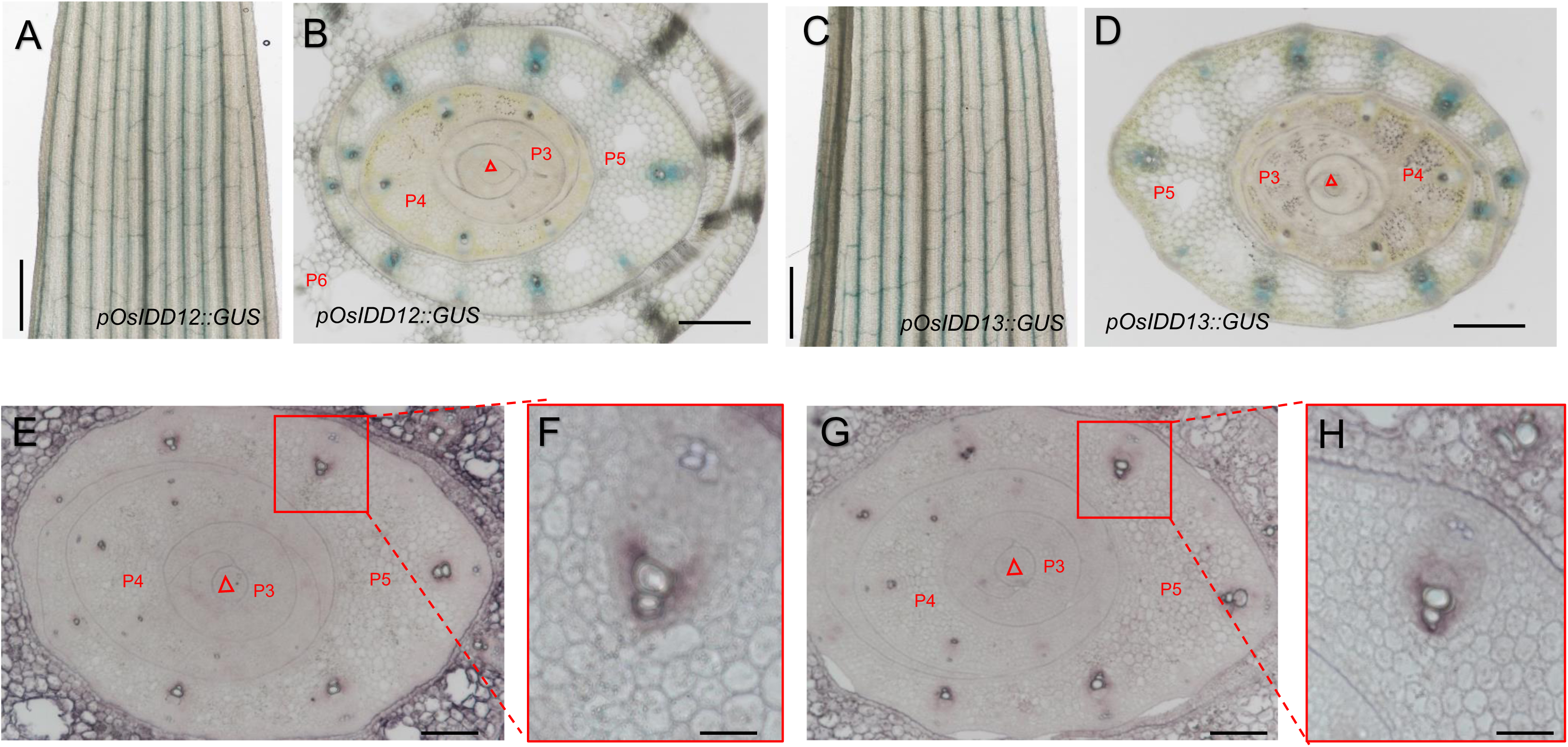
*OsIDD12* and *OsIDD13* expression profiles in leaf tissues. **A-D**, *OsIDD12* and *OsIDD13* expression in vascular files. GUS-stained leaf tissue suggest *OsIDD12* (**A** and **B**) and *OsIDD13* (**C** and **D**) are expressed in vasculature of expanding leaf tips (6th leaf, 20 DAG seedling) and in P3 to P5 leaf primordia (cross sections of 5 DAG seedling), red triangles indicate the position of the SAMs. **E** - **H**, *in situ* analysis using anti-sense probes to *OsIDD12* (**E** and **F**) and *OsIDD13* (**G** and **H**) indicating expression in developing veins of leaf primordia, panels **F** and **H** are close-up images of panel **E** and **G**, respectively showing the specific expression of *OsIDD12* and *OsIDD13* in a major vein of P5. Bars for **A** and **C** = 1 mm, for **B** and **D** = 200 μm, for **E** and **G**= 100 μm, for **F** and **H** = 50.

To test the interaction of IDD proteins with SHR directly, we conducted a yeast two-hybrid (Y2H) screen using OsSHR1 as bait in a screen of a young shoot cDNA library. Of the all clones screened and then sequenced, only OsIDD13 was identified as a putative target (Supplementary Figure 12A). However, given the similarity in sequence identity and expression (Supplementary Figure 11 and Figure 5), we directly tested the interaction of OsSHR1 and OsSHR2 with OsIDD12 and OsIDD13 in Y2H assays (Figure 6A and Supplementary Figure 12B), pull-down assays (Figure 6B and 6C), bimolecular fluorescence complementation assays (BiFC) in rice protoplasts (Figure 6D and 6E) and in tobacco seedling leaves (Supplementary Figure 12C). Collectively, all five assays indicated a specific interaction of OsSHR1 and OsSHR2 with OsIDD12 and OsIDD13.

**Figure 6.**
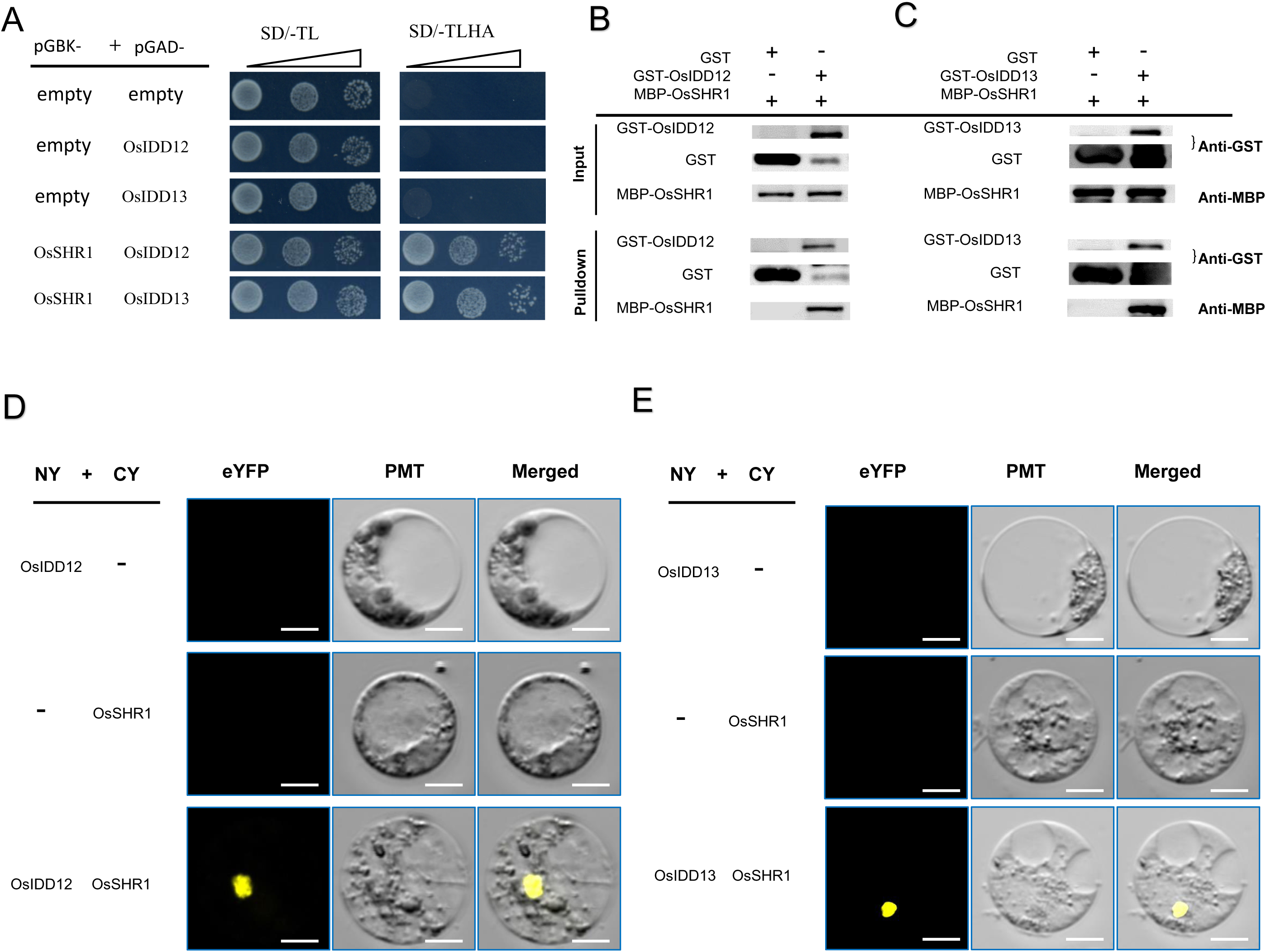
OsSHR1 interacts with OsIDD12 and OsIDD13 directly. **A**, OsSHR1 interacts with OsIDD12 and 13 directly in Y2H assay. OsSHR1 was fused with an in-frame DNA binding domain and OsIDD proteins contained an in-frame activation domain. Transformants were spotted on SD -Leu -Trp -His (TDO) solid media plates with 40 mM 3-AT and plated at 10-, 100-, and 1,000-fold dilutions. **B** and **C**, Pull-down assays revealed rice IDD12 and IDD13 interacting with OsSHR1. The test was performed using OsIDD12 and OsIDD13 in-frame fusions with GST and an OsSHR1 in-frame fusion with the MBP tag. GST: OsIDD12 fusion protein specifically interacted with MBP:OsSHR1 fusion as shown with anti-MBP antibody (**B**), and GST: OsIDD13 specifically interacted with MBP:OsSHR1 analyzed with anti-MBP antibody (**C**). **D** and **E**, BiFC analysis of OsSHR1 interaction with OsIDD12 and OsIDD13 in rice protoplasts, OsIDD12 and OsIDD13 in-frame fusion with N-terminus of split YFP, and OsSHR1 in-frame fusion with C-terminal of split YFP, respectively. Bars for **D** and **E** = 5 μm.

### OsIDD12 and OsIDD13 regulate leaf anatomy in rice

The similarity of expression pattern and direct interactions of rice SHR with OsIDD12 and OsIDD13 strongly suggest that a SHR-SCR complex regulates vascular pattering through the interaction with IDD12 and IDD13. To explore this hypothesis, we ectopically expressed and generated loss-of-function alleles of *OsIDD12* and *OsIDD13* (Supplementary Table 2). OX of *OsIDD12* and *OsIDD13* with native promoters resulted in no significant alteration in M cell number, major vein or minor vein number (Supplementary Figure 13). Similarly, single gene loss-of-function alleles of *OsIDD12* and *OsIDD13* generated with CRISPR-Cas9 (Supplementary Figure 8D) revealed some plant architectural changes, but no significant M cell or vein pattern differences between mutants and WT (Supplementary Figure 14). Thus, we also generated double mutants of *OsIDD12* and *OsIDD13* (*osidd12-3 osidd13-3* and *osidd12-4 osidd13-4*) using CRISPR-Cas9 (Supplementary Figure 8E). Double mutants were short-statured (Figure 7A), with wide leaves (Figure 7B). The number of M cells between two adjacent minor veins decreased while the total number of major and minor veins of double mutants increased significantly (Figure 7C - 7H, Supplementary Figure 15).

**Figure 7.**
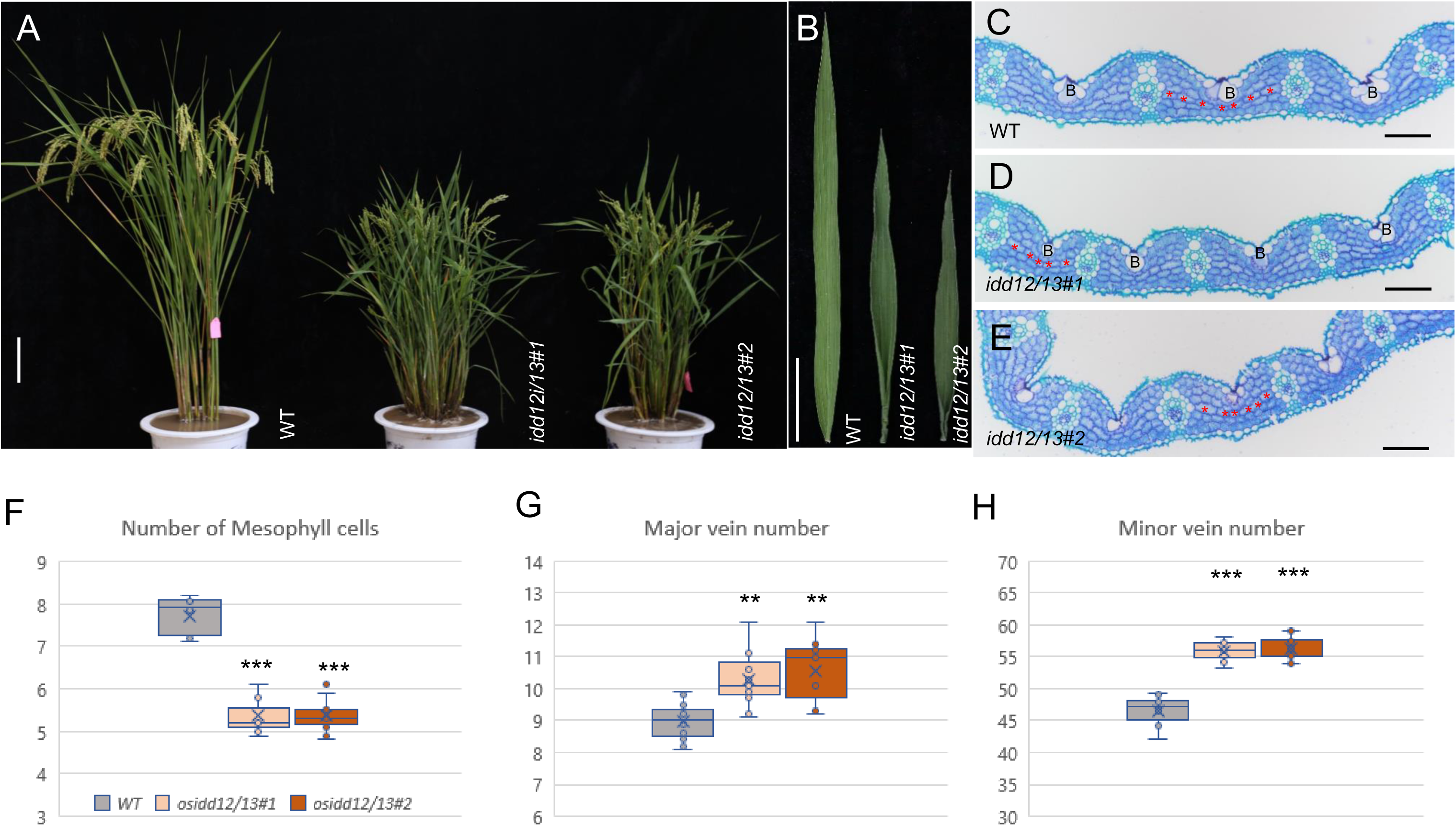
*OsIDD12/13* regulate leaf anatomy in rice. *OsIDD12* (LOC_Os08g36390) and *OsIDD13* (LOC_Os09g27650) double mutants (*idd12/13#1* and *idd12/13#2*) generated by CRISPR-Cas9. **A**, Mature plant images of WT and two independent double mutant lines. **B**, FL of WT and double mutant showing short and wide leaf morphology of mutants. **C**, **D** and **E**, Transverse paraffin sections in mid region of FL showing M cells (red asterisks), B cells and minor veins. Bars for **A** = 20 cm, for **B** = 15 cm, for **C**, **D** and **E** = 100μm, for **F**, **G** and **H**, Leaf trait data in WT and double mutants. P values calculated using student’s t-test, ** represent p<0.01, *** represent p<0.001, n = 11 for each genotype.

### OsIDD12 and OsIDD13 bind a conserved motif in intron 3 of *OsPIN5c*

The PIN family of auxin efflux carriers has long been implicated in the development and differentiation of vascular tissues in the grasses (Forestan and Varotto, 2012; O’Connor et al., 2014, 2017). Thus, we reasoned that the decreased M cell number and increased minor vein number observed in *osidd12 osdd13* double mutants could be due, in part, to a disruption in the expression of *OsPIN* gene family members. A search for putative full-length IDD binding sites in the 12 PIN family members of rice identified a canonical IDD binding site ‘TTTGTCGCTTT’ in the third intron of rice *PIN5c* (Supplementary Figure 16; (Kozaki et al., 2004)) and alternative IDD binding sites in several other *PIN* genes (Supplementary Figure 16; (Seo et al., 2011; Xuan et al., 2012; Sun et al., 2019, 2020)). Preliminary expression analysis of *PIN* family members in the *shr1 shr2* double mutant suggested that expression of *OsPIN5c* may be regulated by SHR. Thus, we focused our molecular characterization on rice *PIN5c* as a potential direct target for IDD binding.

To verify the binding of OsIDD12 and OsIDD13 to intron 3 of *OsPIN5c*, we performed DNA pull-down assays using OsIDD12 and OsIDD13 in-frame fusions with GST as bait and tested binding to fragments of intron 3 DNA (Figure 8A). The DNA fragment containing the ‘ TTTGTCGCTTT ’ motif was highly enriched compared to other regions of intron 3 as detected by qPCR (Figure 8A). Electrophoretic mobility-shift assay (EMSA) assays further suggested that OsIDD12 and OsIDD13 bind the specific ‘TTTGTCGCTTT’ motif (Figure 8B and 8C; Supplementary Figure 17). To verify binding specificity, ChIP-qPCR analyses were performed again showing strong binding to fragments carrying the ‘TTTGTCGCTTT’ motif *in vivo* (Supplementary Table 2 and Figure 8D). Collectively, these data strongly suggest that OsIDD12 and OsIDD13 directly interact with a regulatory element within the *OsPin5c* gene.

**Figure 8.**
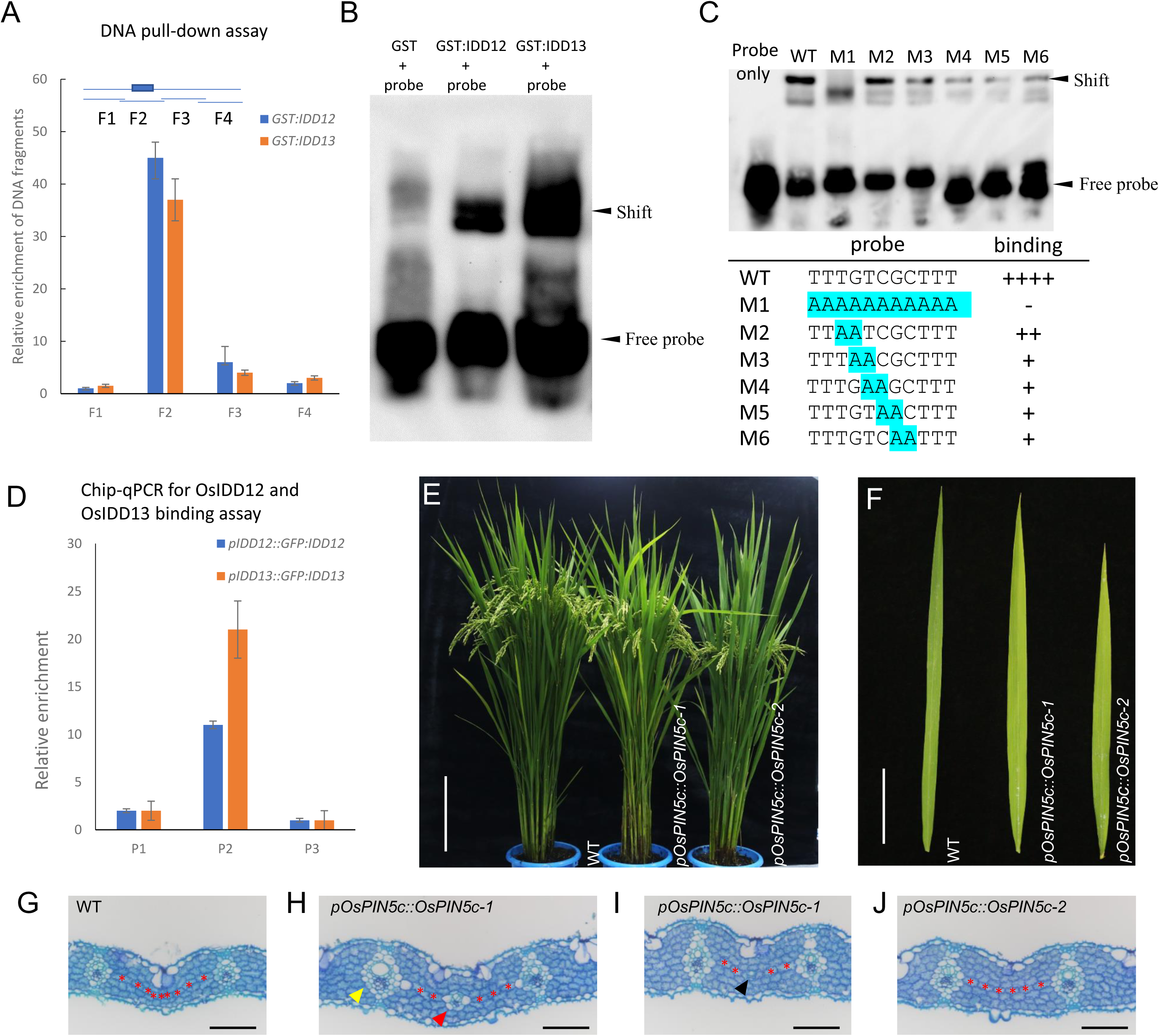
OsIDD12/13 interaction with *OsPIN5c* and the regulation of minor vein initiation by *OsPIN5c*. **A**, OsIDD12 and OsIDD13 in-frame fusions with GST were used as bait to pull down overlapped DNA fragments amplified from the *OsPIN5c* (LOC_Os08g41720) intron 3, input DNA F2 fragment containing putative IDD binding motif (‘TTTGTCGCTTT’) was highly enriched as detected by qPCR (blue box represents IDD binding site in schematic, position of DNA fragments in intron 3 shown by lines). **B**, An electrophoretic mobility-shift assay (EMSA) indicated IDD12 and IDD13 binding the ‘TTTGTCGCTTT’ motif, proteins and probes were equivalently loaded. **C**, EMSA test using different mutant probes for analysis the binding affinity of IDD13, proteins and probes were equivalent loaded. **D**, ChIP-qPCR analysis showing specific binding of IDD with the ‘TTTGTCGCTTT’ motif *in vivo*, assays were performed using the *pIDD12::GFP:IDD12* and *pIDD13::GFP:IDD13* transgenics, qPCR with primers targeted the IDD binding motif (TTTGTCGCTTT) named P2 and two neighboring control DNA fragments named P1 and P3 in the third intron of *OsPIN5c* were tested. **E** and **F**, Images of *OsPIN5c* OX lines showing plants at seed setting stage (**E**) and wider flag leaves (**F**). **G** - **L**, Transverse paraffin sections in mid region of FL including WT and two independent OX lines showing, M cells (asterisks), minor veins defined as rank-1 (**H**, yellow triangle) or rank-2 intermediate veins (**H**, red triangle), and a putative minor vein initial (**I**, black triangle) **J,** typical interveinal pattern showing reduced M cell numbers between minor veins (note reduced bulliform cell development). Bars for **E** = 20 cm, **F** = 15 cm, for **G** - **J** = 100μm.

To directly test the function of OsPIN5c in rice, we ectopically expressed *PIN5c* with both its native and constitutive promoters (Supplementary Table 2). We were unable to generate T_1_ seed when *OsPIN5c* was constitutively expressed (Supplementary Table 2). However, when using the native promoter to drive expression, multiple independent single copy lines were recovered (Supplementary Table 3). Plants displayed similar architecture as wild-type plants (Figure 8E) but with wider leaves (Figure 8F). Interestingly, transverse sections of rice flag leaves of *OsPIN5c* OX lines occasionally conditioned both rank-1 and rank-2 intermediate veins (Figure 8H; as defined by the absence of sclerenchyma cells on the abaxial surface of the vein), and what appear to be vascular initials (Figure 8I). Plants ectopically expressing *PIN5c* also had increased major and minor vein numbers and reduced M cell numbers between minor veins (Figure 8G - 8J and Supplementary Figure 18). Although these phenotypes were not fully penetrant, they nonetheless support the hypothesis that OsIDD12 and OsIDD13 mediate their effect at least partially through an interaction with rice *PIN5c*.

### *OsPIN5c* expression is repressed by OsSHR1/2 and OsIDD12/13

To further explore the potential of *PIN5c* regulation by a SHR/IDD complex, individual leaf primordia were isolated using laser capture microdissection (LCM) and RNA was isolated from P1, P2, P3, P4 and P5 primordia. qPCR was then performed on mRNA isolated from pooled leaf primordia of WT, *SHR* OX lines, *shr* mutant and *idd* mutant lines. Notably, while rice *PIN5c* expression was reduced approximately 50% in *SHR* OX lines (Supplementary Figure 19A) relative to WT, it increased approximately 5-fold in *shr1shr2* double mutants (Supplementary Figure 19B) and over 3-fold in *idd* triple mutants (Supplementary Figure 19C).

### Intron 3 regulates *OsPIN5c* expression in a dual luciferase reporter assay in rice leaf protoplasts cells

We used a dual luciferase (LUC) reporter assay to explore the regulatory network of SHR1/2, IDD12/13, and *PIN5c* (Figure 9). Protoplast cells (PCs) from two-week-old Nipponbare seedlings were used to introduce expression constructs (Figure 9A). As expected, control vectors carrying the HA epitope tag alone (*p35S::HA)* or HA fused to OsIDD12 (*p35S::OsIDD12:HA)* did not induce luciferase from the promoter-less pLUC construct (Figure 9B). When the *PIN5c* promoter was inserted upstream of the luciferase reporter (*pProPIN5c::LUC*), robust luciferase expression was observed, indicating that the *PIN5c* promoter fragment can drive gene expression in this protoplast reporter system. When IDD12 was introduced (*p35S::OsIDD12:HA),* LUC expression did not increase above the *35S* control line suggesting that IDD12 does not directly inhibit gene expression from the *PIN5c* promoter. However, when the intron 3 fragment carrying the IDD binding-motif ‘TTTGTCGCTTT’ was inserted upstream of the *PIN5c* promoter (*pI3-ProPIN5c::LUC*), relative LUC activity was reduced by about half compared to that *ProPIN5::LUC* suggested the *cis*-element ‘TTTGTCGCTTT’ is capable of mediating *OsPIN5c* repression. Introduction of *p35S::OsIDD12:HA* together with the *pI3-ProPIN5c::LUC* reporter, failed to result in further reductions in luciferase expression. Similar results were obtained when a construct carrying OsIDD13 was used in the dual luciferase assays (Figure 9C). Namely, that OsIDD13 did not increase repression mediated by the intron 3 fragment in a wild-type Nipponbare background. These results are consistent with the finding that overexpression and loss-of-function alleles of *IDD12* or *IDD13* did not affect *PIN5c* mRNA accumulation (Supplementary Figure 19C and 19D) or leaf anatomy (Supplementary Figure 14) and suggests that IDD12 and IDD13 may act cooperatively to mediate repression of *PIN5c* by SHR.

**Figure 9.**
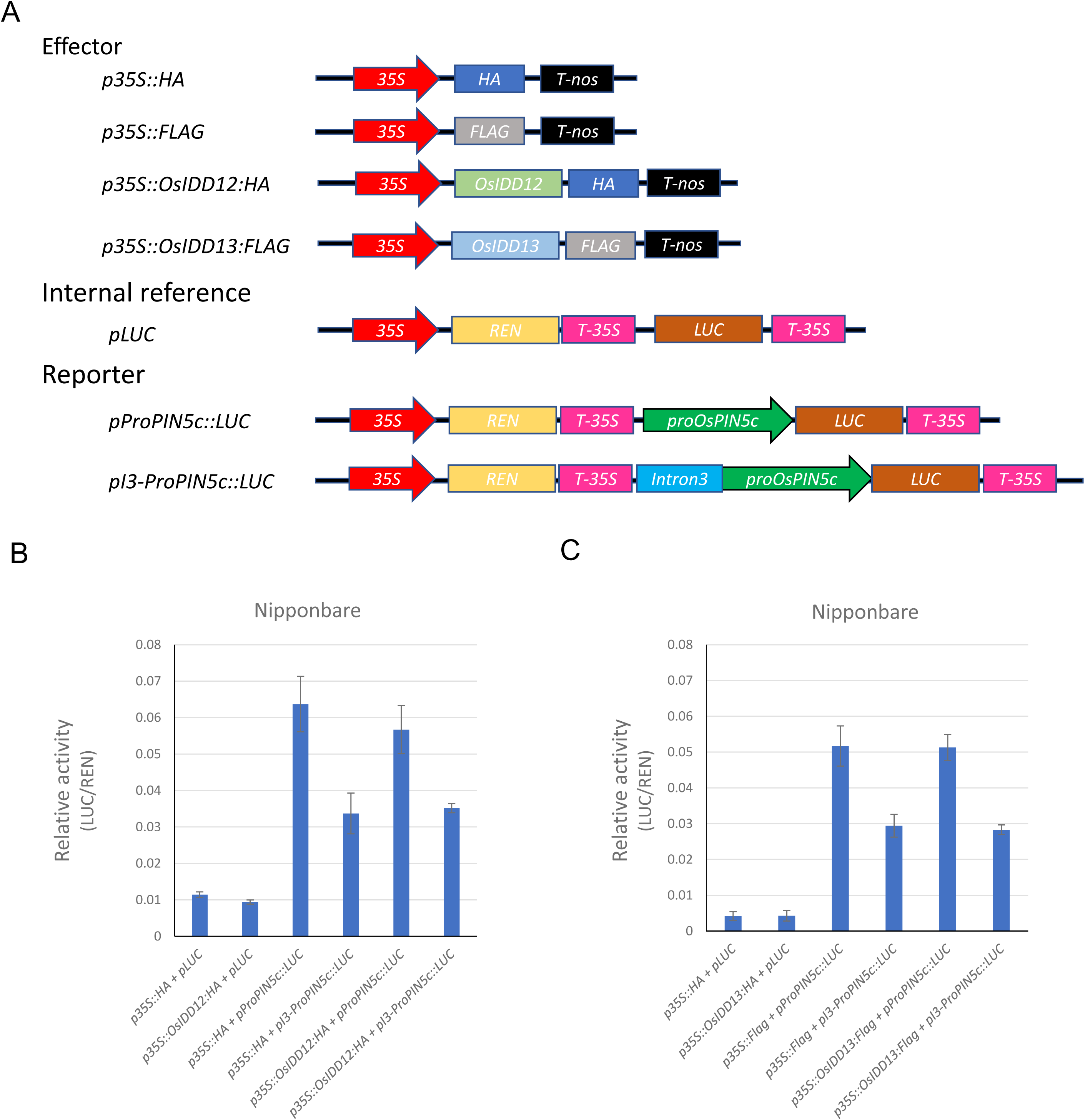
*OsIDD12* and *OsIDD13* do not directly regulate *OsPIN5c* expression in rice leaf protoplasts cells. Rice leaf protoplasts cells isolated from two week old seedlings. LUC relative activity is shown as a readout of transcriptional activity and three biological replicates were performed for all assays. **A**, Schematic diagrams of the reporter and effector constructs. The firefly luciferase (LUC) gene driven by the *OsPIN5c* promoter and the *Renilla* luciferase (REN) reporter gene driven by the 35S promoter were used as the reporter and internal controls, respectively. For the effectors, OsIDD12 and OsIDD13 were fused with a HA or FLAG tag, respectively. **B** and **C**, Constructs carrying *IDD12* (**B**) or *IDD13* (**C**) with the third intron (I3) plus the *OsPIN5c* promoter region (*pI3-ProPIN5c* fusion) driving *LUC* gene were introduced into WT protoplasts. Neither IDD12, nor IDD13 enhanced inhibition of *OsPIN5c* expression.

### Ectopic expression of *OsPIN5c* rescues *SHR* overexpression phenotypes

To genetically test the hypothesis that SHR directly mediates the repression of *PIN5c* in rice, we introduced *PIN5c* OE constructs into *SHR1* overexpression lines with the expectation that expression of a *PIN5c* cDNA clone lacking the IDD binding motif would overcome *SHR* repression. Callus of transgenic line *pOsSHR1::OsSHR1-1* was retransformed with the *OsPIN5c* cDNA driven by its native promoter (Supplementary Table 2) and T_0_ lines screened for single copy *pOsPIN5c::OsPIN5c* insertions (Supplementary Table 3). Two lines were selected and self-pollinated to generate segregating T_1_ plants for detailed analysis (Figure 10A). Approximately 1/4 of the plants segregated for a rolled leaf phenotype while 3/4 had wild-type-like, flat leaves, consistent with a rescue of *SHR* OE leaf rolling phenotype by ectopic expression of *OsPIN5c* (line 1=17 flat leaf plants : 6 rolled leaf plants = 2.83 : 1, x^2^= 0.0145; line 2 = 22 flat leaves plants : 7 rolled leaf plants =3.14 : 1, x^2^= 0.0142). To confirm that the phenotypic effects were not due to silencing of the *SHR* transgene, *SHR1* and *PIN5c* gene expression was measured in the segregating lines, indicating that the rescue of the leaf rolling phenotype was mediated by ectopic *PIN5C* expression (Figure 10B). Characterization of additional leaf phenotypes including M cell number between veins, minor vein number and bulliform cell area are consistent with the *PIN5c* OE phenotypes displayed in WT plants (see Figure 8; Supplementary Figure 18) and suggests that in the absence of the IDD binding motif, SHR cannot mediate repression of *OsPIN5c* even when overexpressed.

**Figure 10.**
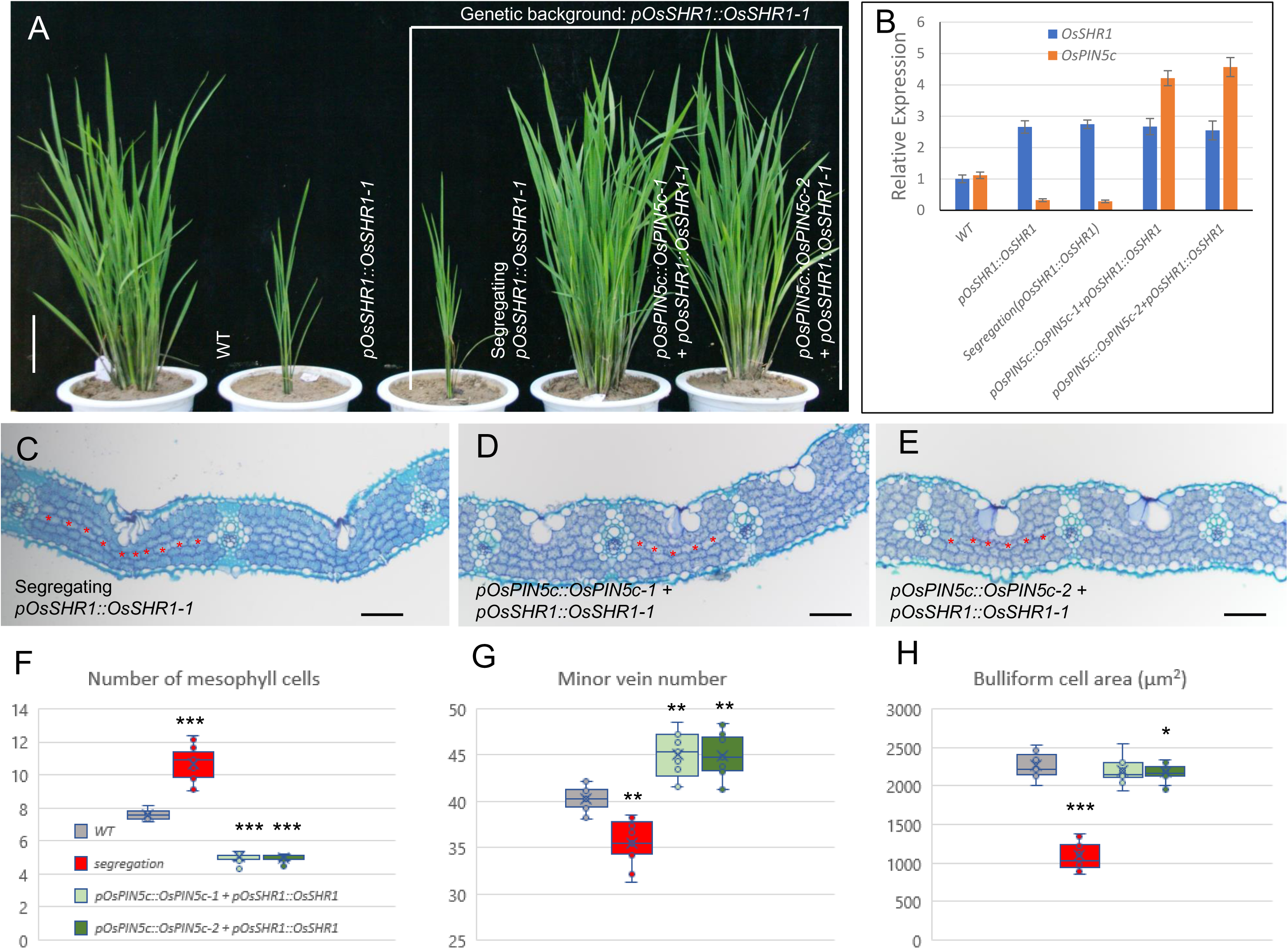
*OsPIN5c* rescues *SHR* overexpression phenotypes. **A**, Images of WT, homozygous *OsSHR1-1* OX plant, segregating plant with *SHR1* OX morphology, and two independent T_2_ *OsPIN5c* OX plants that also carry the *pOsSHR1::OsSHR1* transgenes (OX-1: *pOsPIN5c::OsPIN5c-1* + *pOsSHR1::OsSHR1-1* and OX-2: *pOsPIN5c::OsPIN5c-2* + *pOsSHR1::OsSHR1-1*). **B**, Relative expression of *OsSHR1* and *OsPIN5c* in WT, *SHR1* OX plant and *PIN5c* OX plants in *SHR1* OX background. **C** - **E**, Transverse paraffin sections in mid region of leaves showing M cells, B cells and minor veins including WT (**C**), and OX-1 (**D,** *pOsPIN5c::OsPIN5c-1* + *pOsSHR1::OsSHR1-1*) and OX-2 (**E**, *pOsPIN5c::OsPIN5c-2* + *pOsSHR1::OsSHR1-1*) plants. **F**, **G** and **H**, Leaf trait data in WT and segregating lines of indicated genotypes. P values calculated using student’s t-test, * p<0.05, ** p<0.01, *** p<0.001, n = 11 for each genotype. Bars for **A** = 20 cm, for **C**, **D** and **E** = 100μm.

### *idd12*/*13* double mutants are epistatic to SHR overexpression

*idd12*/*13* double mutants and *SHR* single or double mutants have fewer M cells between minor veins and have increased minor vein number (See Figures 2 and 7). The direct interaction of SHR with IDD12/13 further suggests that SHR and IDD regulate downstream target genes through complex formation. To test this hypothesis, *SHR1* and *SHR2* were overexpressed in an *idd12*/*13* double mutant background (*osidd12-4 osidd13-4*). Introduction of *SHR1* and *SHR2* OX constructs into *osidd12-4 osidd13-4* and the increased expression of *SHR1* or *SHR2* (supplementary Figure 20A and 20B) had no significant effect on M cell number or minor vein numbers relative to the *idd12 idd13* mutant background (Figure 11A), consistent with this hypothesis.

**Figure 11.**
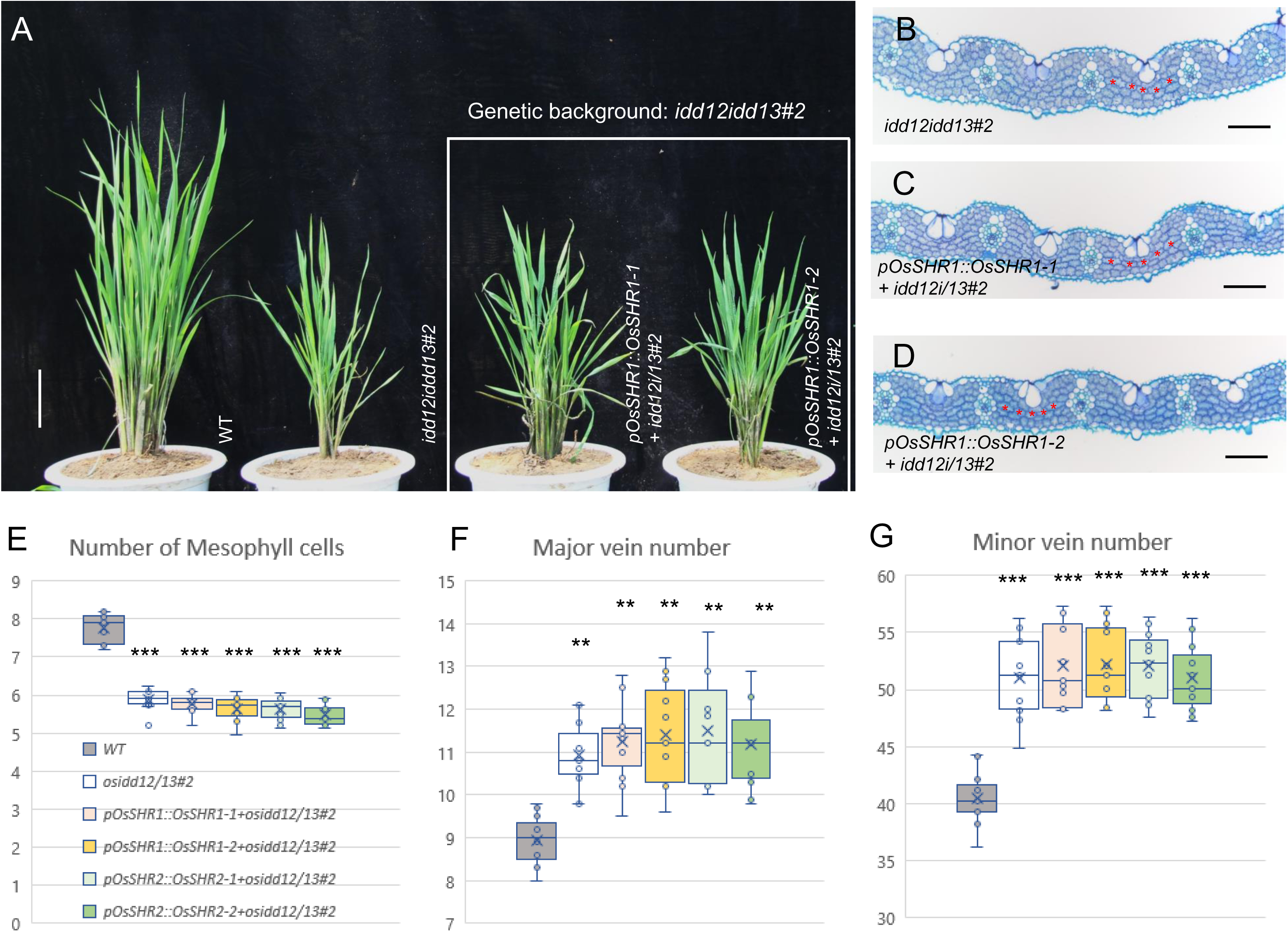
*idd12/13* double mutant phenotypes are epistatic to SHR overexpression. T_2_ *idd12/idd13* double mutant (*idd12/13#2*) used as donor material for genetic transformation of *OsSHR1* OX lines driven by its native promoter, panel **A** showing phenotype of 12 week old seedlings. **B**, **C** and **D**, Transverse paraffin sections in mid region of leaves showing M cells, B cells and minor veins of *idd12/13#2* (**B**), OX-1 (**C**, *pOsSHR1::OsSHR1-1 + idd12i/13#2*) and OX-2 (**D**, *pOsSHR1::OsSHR1-2 + idd12i/13#2*) plants. **E**, **F** and **G**, Leaf trait data of indicated genotypes. P values calculated using student’s t-test,** p<0.01, *** p<0.001, n = 11 for each genotype. Bars for **A** = 20 cm, for **B**, **C** and **D** = 100μm.

### OsSHR1, OsSHR2, OsIDD12 and OsIDD13 regulate *OsPIN5c* expression in a dual luciferase reporter assay

To directly test the roles of SHR1 and SHR2 in the transcriptional regulation of *PIN5c*, we isolated protoplasts from the *shr1 shr2* double mutant for the dual luciferase assay. As shown in Supplementary Figure 21A and 21B, we no longer observed the reduced expression of the *LUC* gene mediated by the intron 3 fragment (see also Figure 9) in the double mutant background indicating that SHR is necessary for *PIN5c* transcriptional repression. Furthermore, when protoplasts were isolated from *SHR1* or *SHR2* OX plants, intron 3 - mediated repression of *OsPIN5c* was dramatically enhanced conditioning approximately 10% of the relative activity of the wild-type *PIN5c* promoter (Supplementary Figure 21C and 21D). When protoplasts were isolated from *idd12 idd13* mutant plants we also failed to see repression of *PIN5c* mediated by intron 3 (Supplementary Figure 22A, 22B and 22C). Interestingly, when protoplasts constitutively expressing IDD13 were used for transfection (Supplementary Figure 22D), IDD12-mediated repression of LUC expression in the presence of intron 3 was greatly enhanced. Similar results were observed when protoplasts expressing IDD12 were transfected with IDD13 (Supplementary Figure 22E) suggesting that IDD12 and IDD13 act cooperatively to bind to the intron 3 motif. In summary, the results of genetic, *in vitro* and *in vivo* protoplast assays all support the hypothesis that SHR repression of *OsPIN5c* expression is dependent on both IDD12 and IDD13 binding to intron 3 of *OsPIN5c*.

## Discussion

### SHR promotes ground cell fate in grass leaves and inhibits minor vein formation

Previous attempts to interpret the function of SHR by ectopic overexpression with strong constitutive promoters in rice have largely been unsuccessful resulting in few if any recovered plants (Wu et al., 2014; Schuler et al., 2018), whereas the use of weak promoters to drive SHR expression suggested a limited role of SHR in grass leaf development (Schuler et al., 2018). We also observed that in the course of generating over 200 independent T_0_ lines carrying constructs driving *OsSHR1*, *OsSHR2* and maize *SHRs* (*ZmSHR1*, *ZmSHR2/2h*) with *35S* or actin promoters, only a handful of plants survived beyond the young seedling stage and even fewer produced viable seeds (see Supplementary Table 2 and Supplementary Figure 2). These results suggest that high levels of SHR likely lead to severe disruptions in multiple cellular identities, including those in reproductive organs. Thus, for the majority of our analyses we focused on the characterization of lines generated using endogenous promoters of *SHR1* and *SHR2* to maintain tissue-specific controls but to moderately enhance expression. Nevertheless, we observed dramatic phenotypes in leaf tissues when either *SHR1* or *SHR2* were expressed in vascular tissues. A recent study of ectopically expressed *ZmSHR1* in maize revealed ZmSHR1 is capable of traversing at least 8 cell layers of the root cortex (Ortiz-Ramírez et al., 2021). Given the restricted expression of *SHR1* and *SHR2* to vascular tissues in the leaf (Figure 4; (Schuler et al., 2018)), our data suggests that SHR protein is also capable of trafficking multiple cell layers in leaf tissue and may directly mediate the proliferation of M cells much as it mediates cortex differentiation in the root.

A high vein density is a structural characteristic of nearly all C_4_ leaves (Sedelnikova et al., 2018), and is considered a necessary prerequisite to support the metabolic cooperation of a two-cell C_4_ photosynthetic system (Hibberd et al., 2008). As was observed in previous studies of maize (Slewinski et al., 2014), disruption of SHR function in *S.viridis* alters Kranz anatomy resulting in higher vein densities at the expense of M cell numbers. A similar increase in vein density was also observed in *scr1/1h* double mutants in maize (Hughes et al., 2019), suggesting that both SCR and SHR regulate the ratio of ground to vascular cells in C_4_ grasses and play either a direct or indirect role in mediating BS and M cell identities (Slewinski et al.,2013, 2014; Hughes et al., 2019). However, it is worth noting that neither triple mutants of *shr* (*Zmshr1*;*Zmshr2*;*Zmshr2h*) nor triple mutants of *scr* (*Zmscr1*;*Zmscr1h*;*Zmscr2*) have been characterized in maize. Given the redundant roles of SHR1 and SHR2 observed here in rice leaf development, it is likely that these higher order mutants would reveal much more about the SHR-SCR regulatory network than can be deduced from single or even double mutant studies. Although double mutants of *svshr1 svshr2* have recently been reported (Ortiz-Ramírez et al., 2021), characterization of the leaf phenotypes conditioned by these double mutants has not yet been described and we were unable to generate the *svshr1 svshr2* double mutants in our study.

In rice, the role of SHR in leaf development could be more directly tested through the characterization of *osshr1 osshr2* double mutants. Consistent with the observations in maize and *Setaria* detailed above, OsSHR1 and OsSHR2 act redundantly to promote M cell identity and determine minor vein placement. However, in rice, SHR1 and SHR2 also repress the differentiation of bulliform cells and promote sclerenchyma cell development on the abaxial surface of minor veins. We also observe aberrant stomatal placements on both adaxial and abaxial surfaces of the leaf, but fail to detect *SHR* expression in epidermal cells, suggesting that the protein is capable of modulating epidermal cell fates through a mobile signal.

### IDD12 and IDD13 interact with SHR to regulate vein patterning in rice

Several studies have indicated that IDD proteins interact with SHR in *Arabidopsis* or rice including a crystal structure of the IDD protein complexed with SHR (Welch et al., 2007; Hirano et al., 2017). OsIDD13, together with OsIDD3 and OsIDD14/OsLPA1 have been reported to likely form a transcriptional complex that directly regulates *PIN1a* in rice (Sun et al., 2019, 2020). In *A. thaliana*, *IDD14*, *IDD15*, and *IDD16* are expressed in vascular cells and act redundantly to regulate auxin biosynthesis through the direct interaction with promoters of several auxin pathway genes (Cui et al., 2013). We found ectopic expression of either rice *IDD12* or *IDD13* alone did not affect the anatomy of leaves suggesting that the stoichiometry of the IDD and SHR proteins is important for their function. Importantly, the *osidd12 osidd13* double mutants did condition reduced M cell numbers and increased minor veins, phenotypes that are consistent with *SHR* double mutants, providing the strongest genetic evidence to date that SHRs and IDDs together regulate grass leaf anatomy.

This genetic evidence was further supported by the *in vitro* Y2H, pull-down and BiFC assays demonstrated that rice SHR and IDD12/13 physically interact as well.

### A central role for *PIN5c* in mediating the SHR-IDD regulation of vein formation

The results of *in situ* analysis suggests that in WT, *OsSHR1* is restricted to cells within vascular files and we speculate that this may result in a restricted auxin domain through the decreased expression of a subset of PIN proteins. As previous reported, rice IDD proteins are capable of directly binding to the *PIN1a* promoter (Sun et al., 2019, 2020), thus we screened all the rice PIN gene sequences including promoters, exons and introns for both canonical and non-canonical IDD binding motifs (Kozaki et al., 2004; Kobayashi et al., 2017) and identified a canonical binding site in the third intron of *PIN5c*. DNA binding, EMSA and ChIP-qPCR assays verified the physical interaction of OsIDD12 and OsIDD13 to the ‘TTTGTCGCTTT’ motif. To validate this finding, we further conducted a series of transcriptional assays using the dual luciferase reporter system (De Sutter et al., 2005; Grunewald et al., 2012; Pollier et al., 2013; Liu et al., 2014; Cárdenas et al., 2016). These studies supported the results of genetic and molecular interaction assays and further showed that the IDD binding motif is sufficient to mediate the SHR and IDD-dependent repression of *PIN5c* driven expression. A model describing these interactions is provided in Figure 12. Ongoing ChIP-seq and ATACseq studies could provide further insights into the global regulatory networks mediated by the SHR and IDD family members in the grasses.

**Figure 12.**
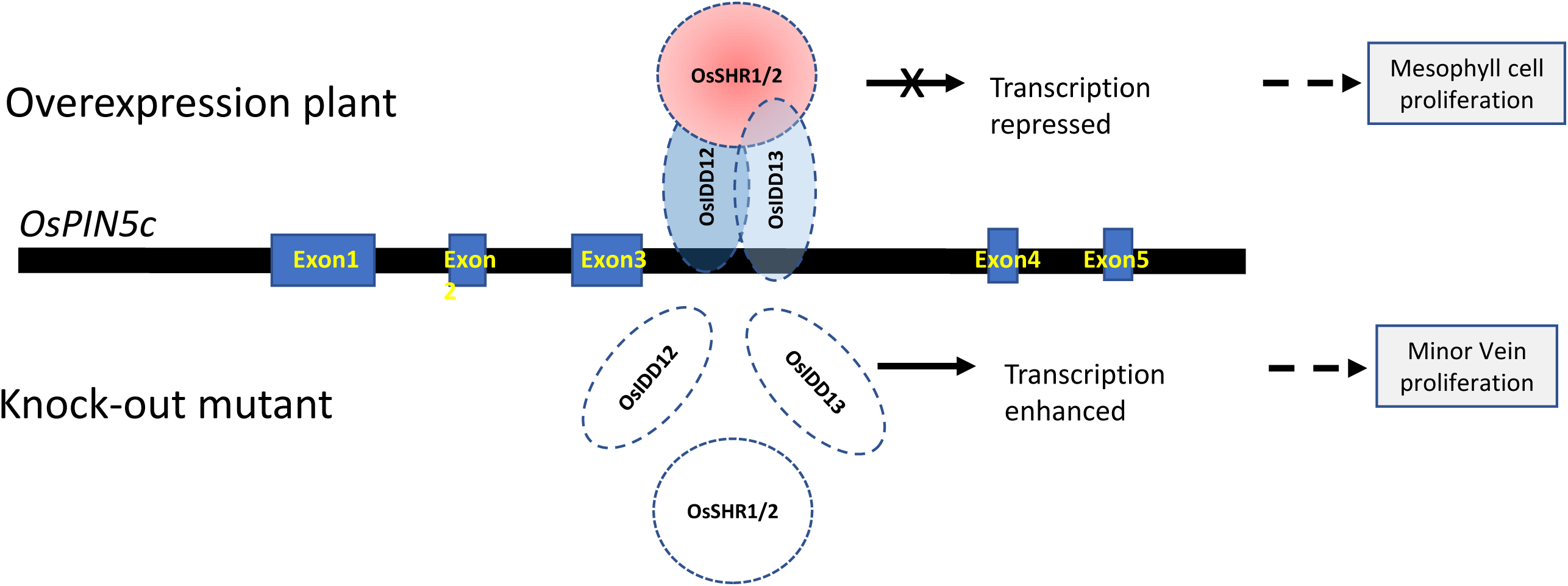
Working model for *OsPIN5c* regulation by OsSHR1/2 and OsIDD12/13. Working model of rice *SHR/IDD* regulation of *OsPIN5c* expression. In OX lines (upper panel), increased expression of OsSHRs proteins enhance OsIDDs binding to the ‘TTTGTCGCTTT’ motif present in intron 3 of *PIN5c* resulting in repression of transcription from the gene and promoting ground cell development (see supplemental figure 19**A**). Conversely, loss-of-function alleles of IDD or SHR (lower panel) result in the de-repression of *PIN5c* rice (see supplemental figure 19**B** and supplemental figure 19**C**) and the proliferation of minor veins.

### Optimized manipulation of the *SHR*-*IDD*-*PIN5c* circuit as a foundation for Kranz-like anatomy in C_3_ grasses

As noted earlier, a primary objective of engineering C_4_ traits into C_3_ grasses relies on the installation of a C_4_ vascular chassis to support C_4_ biochemistry. Importantly, the mutant characterization presented for *shr* and *idd* mutants reveal several traits that support a C_4_ anatomy including increased vein density, appearance of morphologically distinct minor veins, reduced M cell numbers, reduced lobing of M cells, and thickened cell walls of vascular bundles. Nevertheless, the pleiotropic nature of the SHR and IDD mutant phenotypes poses significant challenges including shorter statured plants, low yield and reduced gravitropic response. In contrast, rice *PIN5c* OX plants displayed relatively normal growth habit and good fecundity. Thus, we suggest, that further manipulation of the rice *PIN5c*, perhaps in combination of weaker alleles of the *IDD* genes presents an ideal target for optimizing anatomical traits in rice to support a C_4_ biochemistry.

## Methods

### Plants materials and growth conditions

Rice (*Oryza sativa* L.) japonica cultivar ‘Nipponbare’ and *Setaria viridis* accession ME034V were used as plant material for experiments. All mature rice plant phenotypic characterizations were performed on plants grown in paddy fields under natural light conditions at the Chinese Academy of Agricultural Sciences (CAAS, Beijing, China) transgenic field facility (Langfang, Hebei, China). Rice seedling tissues for GUS expression analysis, *in situ*, BiFC, Chip-qPCR, root growth assays, LCM, Dual luciferase reporter assay and qPCR analysis were grown in growth chambers (AR-36L2, Percival Scientific, USA.) with day/night temperature maintained at 28℃ /22 ℃ and a diurnal light regime of 16 hours of light and 8 hours of dark at Biotechnology Research Institute (BRI, CAAS). Seeds were propagated at both the Langfang facility from May to October and in the paddy field under natural growth conditions at the CAAS field station in Sanya (Hainan, China) from November to March. *S. viridis* ME034V plants were grown in growth chambers or greenhouses at BRI with conditions as described (Jinek et al., 2012). Briefly, seedlings were grown under similar conditions at 12h day/12h night regime with 50% humidity and 31℃/22℃ (day/night) temperature.

### Generation of binary constructs and plant transformation

The binary vector *pCSpeGH* (NCBI accession number: MW186668) was modified from vector *pCambia1391Z* (Hajdukiewicz et al., 1994) to generate OX constructs. Firstly, *pCambia1391Z* was digested with *Sac*II (TAKARA, Japan), then ligated with a *SpeI* DNA fragment amplified from vector *pCHF3* (kindly provided by Zhihong Lang, BRI) that carried spectinomycin antibiotic resistance in bacteria to generate the intermediate step plasmid *pCSpe*. Second, to facilitate cloning, we introduced a Gateway-compatible cloning system. Vector *pCSpe* was digested with *Hind*III (TAKARA) and *Pmac*I (TAKARA) and a Gateway^®^ cassette amplified from vector *pGWB14* (kindly provided by Pinghua Li, Shandong Agricultural University, Shandong, China) was ligated using the In-Fusion kit (Clontech, USA) for generation the binary destination vector *pCSpeGH* (Sequence deposited in NCBI Gene Bank with the accession number MW186668). Third, the open reading frames (ORF) of *OsSHR1*, *OsSHR2*, *OsIDD12*, *OsIDD13* and *OsPIN5c* were amplified from cDNA of Nipponbare using GATEWAY compatible primer pairs (Supplementary Table 4). Then, the amplified products were subcloned in the Gateway donor vector *pDONR207* (Invitrogen, USA) by a BP reaction and sequenced. The resultant entry clones were cloned into *pCSpeGH* by LR reaction to generate the destination vectors. Finally, destination vectors were digested and ligated to the proximate 2.5kb promoter fragment of *OsSHR1*, *OsSHR2*, *OsIDD12*, *OsIDD13* and *OsPIN5c* amplified from rice genomic DNA (primer pairs listed in Supplementary Table 4) or to the rice *Actin1* promoter (amplified from *pCam23A*, (Acharya et al., 2017)) or to the CaMV *35S* promoter (amplified from vector pTCK309 (Sun et al., 2016)) respectively, to generate the final vectors for overexpression of the selected genes with native or constitutive promoters.

The promoter fragments of *OsSHR1*, *OsSHR2*, *OsIDD12*, *OsIDD13* and *OsPIN5c* described above were also amplified and inserted into the *pCambia1391Z* (Hajdukiewicz et al., 1994) vector to generate the corresponding promoter::GUS vectors for gene-specific tissue expression analysis.

The binary vectors for ChIP-qPCR analysis were constructed by first amplifying *OsIDD12*, *OsIDD13* cDNA fragments along with *GFP* fragments. Using an overlapping PCR method, we generated the *GPF*:*OsIDD12* and *GPF*:*OsIDD13* PCR fusion proteins (primer pairs listed in Supplementary Table 4), that were then subcloned into *pDONR207* by BP reaction and sequenced. Second, the resultant entry clones were cloned into *pCSpeGH* through an LR reaction to generate the final vectors. Finally, all the *IDD* native promoter sequences were amplified and inserted into the final vectors as described above to generate the binary vectors for Chip-qPCR analysis.

The *OsSHR1*, *OsSHR2*, *OsIDD12*, and *OsIDD13* mutant constructs utilized the CRISPR-Cas9 system (kindly provided by Yao-Gung Liu (Ma et al., 2015)), and two gRNAs were designed to each gene to increase the frequency of deletions. The gRNAs were designed to target the *OsSHR1*, *OsSHR2*, *OsIDD12* and *OsIDD13* coding sequences and assembled into the final binary CRISPR vectors, primers used for constructs are listed in Supplementary Table 4.

The binary vector *pCSpeGH* was first modified for efficient transformation of ME034V. In brief, the promoter of transgenic selection marker cassettes (*35S* drives hygromycin phosphotransferase (HPT) was replaced with the *Ubi* promoter cloned from Switchgrass (*Panicum virgatum*) and the new construct named *pCSpeGHu* (primer pairs listed in Supplementary Table 4). The full-length codling ORFs with stop codons of *SvSHR1*(Sevir.2G383300) and *SvSHR2* (Sevir.9G361300) were amplified from ME034V using gateway compatible primer pairs (Supplementary Table 4), and the final binary vectors driven by native and constitutive promoters and for transformation of *Setaria* were constructed similarly to those described above.

To generate mutants in *SvSHR1* and *SvSHR2*, CRISPR-Cas9 gene editing technology was used (*pJG471*, *pJG310* and *pJG338* vectors carrying CRISPR-Cas9 were kindly provided by Pinghua Li, Shandong Agricultural University). Guide RNAs (gRNAs) were designed to generate single or double mutants of *SHR*, and the primers are listed in Supplementary Table 4.

Constructs for overexpression of *OsPIN5c* or *OsSHR1*/*2* were based on vector *pCam23A* (Acharya et al., 2017). In brief, *OsSHR1*/*2* or *OsPIN5c* cassettes were amplified from *pOsSHR1::OsSHR1*, *pOsSHR2::OsSHR2* or *pOsPIN5c::OsPIN5c* and inserted into the *Kpn*I digested vector *pCam23A* (Sun et al., 2016) backbone by using the In-Fusion kit (Clontech), with primers are listed in Supplementary Table 4.

All the above vectors were sequenced to verify the integrity and orientation of the insertions. The binary constructs were introduced into the *Agrobacterium tumefaciens* strain AGL1 by electroporation. For rice transformation, a tissue culture-based method was used to introduce the binary constructs into rice explants as previously described (Wan et al., 2009). For *Setaria* transformation, as described in (Brutnell et al., 2010), callus was initiated from mature embryos of sterilized *S. viridis* ME034V seeds, incubated with *Agrobacterium* suspension, and selected with 40 mg/l hygromycin. Callus was subsequently transferred to plant regenerative medium to grow shoots and then to rooting generating medium to recover roots under constant selection of 20 mg/l hygromycin.

### DNA and RNA extractions

Genomic DNA for genotyping was isolated from Nipponbare or ME034V using a modified 2% (v/v) cetyl-trimethyl-ammonium bromide (CTAB) method (Murray and Thompson, 1980). High quality rice or ME034V genomic DNA was isolated for TagMan q-PCR or gene/promoter amplification using the DNeasy Plant Mini Kit (QIAGEN, Germany). Rice or ME034V RNA was extracted from various tissues using TRIzol reagent (Invitrogen, USA), the RNA from LCM isolated tissues was extracted using the isolation kit EASYspin micro kit (Aidlab Biotechnologies, Beijing, China).

### Protein expression, extraction and Western blotting

The MBP-OsSHR1, MBP-OsSHR1, GST-OsIDD12 and GST-OsIDD13 fusion proteins were expressed in *Escherichia coli* BL21(DE3) Rosetta strain (Novagen, USA). Frozen cells were extracted with 100 μl of extraction buffer containing 50 mM Tris-HCl (pH 7.6), 15 mM MgCl2, 0.1 M KCl, 0.25 M sucrose, 10% glycerol, 1 mM PMSF (phenylmethylsulfonyl fluoride), protease inhibitor mixture (Sigma-Aldrich, USA), and 14 mM β-mercaptoethanol. After centrifugation of the sample at 15,000 x g for 10 min, the supernatant was sampled, and the amount of protein for each extract was measured according to the manufacturer’s instructions (Bio-Rad, USA). Proteins were detected using anti-MBP (Abcam, USA) or anti-GST antibodies (Abcam) antibodies at proper dilution and visualized with enhanced chemiluminescence (ECL) reagent (GE Healthcare, USA).

Total proteins were extracted from 20 DAG rice seedlings in RIPA buffer containing 50 mM Tris-HCl (pH 7.2), 150 mM NaCl, 10 mM MgCl2, and 1 mM PMSF, protease inhibitor mixture (Sigma-Aldrich, USA), and the supernatant was collected. The total protein concentration was determined using a Bradford protein assay kit (Bio-Rad). Proteins were detected using anti-GFP (Abcam) antibodies at proper dilution and visualized with an ECL reagent (GE Healthcare).

### TaqMAN assay to estimate transgene copy number

Transgene copy number was determined in overexpression lines in T1 and T2 generations using a TaqMAN qPCR assay as previously described (Yang et al., 2005). We used the rice endogenous reference gene *OsRBCS1* coding Rubisco small subunit 1 as a single copy reference gene and determined the copy number of both the *HptII*, *NptII* and rice/*Setaria SHRs*, *IDDs* and *PIN5c* genes in transgenic rice plants. Primer pairs and probes for real-time quantitative PCR are described in Supplementary Table 4. Primers and probes were labeled with the fluorescent reporter dye 6-carboxy-fluoroscein (FAM) on the 5’ end and the fluorescent quencher dye 6-carboxytetramethylrhodamine (TAMRA) on the 3’end. The following probes and amplicon length are shown in Supplemental Table 4 and confirmed samples were used for phenotypic characterizations.

### Histology and microscopy

Light microscopy used to examine leaf anatomy, and tissue was first fixed in formalin– acetic acid-alcohol (FAA) for two rounds of at least 24 hours at room temperature, softened with a 20%(v/v) hydrofluoric acid bath for at least 3 days, and dehydrated through a graded ethanol series (75%-85%-95%-100%-100%) for one to two hours at each ethanol concentration. Samples were embedded in paraffin (Sigma-Aldrich) and 8μm sections were cut with a microtome (RM2155, Leica, Germany) and stained with a filtered 1% toluidine blue or 1% fast green solution (Sigma-Aldrich). Images were taken with an ICC50 HD microscope (Leica)

### Electron microscopy

To analyze flag leaf ultrastructure, transmission electron microscopy (TEM) was performed. Rice leaf segments were dissected by hand with a razor blade and immediately placed into fixation buffer (5% glutaraldehyde, 50 mM sodium cacodylate buffer, pH 7.2). Samples were then immediately vacuum infiltrated to fix the samples. After 24 hours fixation, tissues were rinsed three times for 40 s each in a buffer solution (50 mM sodium cacodylate, pH 7.2). Samples were then transferred to a 1% osmium tetroxide solution for 4 min, and rinsed with water for 40 s. A graded acetone series (30%-50%-70%-80%-90%-95%-100%-100%-100%) was used to dehydrate the samples (one hour at each concentration) which were then infiltrated with resin. Ultrathin sections of approximately 100nm were cut on an ultramicrotome, lifted onto glow-discharged carbon-coated copper grids stained with 0.5% formvar, and observed at 80 kV on a TEM.

For scanning electron microscopy (SEM), flag leaf segments were dissected by hand with a razor blade and immediately placed into fixation buffer. After 24 hours fixation, the samples were dried in a vacuum desiccator for 10 days, coated with gold particles in an argon sputter coater and visualized with a Zeiss Leo 1430 VP SEM.

### GUS expression analysis

*OsSHR1*, *OsSHR2*, *OsIDD12*, *OsIDD13* and *OsPIN5c* promoter::GUS vectors and the control vector *pCAMBIA1391Z* (Hajdukiewicz et al., 1994) were transformed into WT plants, and at least 20 independent transgenic events were assayed for promoter expression. Three single copy lines (Supplementary Table 4) displayed the same GUS activity and patterns chosen for final analysis. Various tissues of T_2_ promoter::GUS plants were incubated in staining solution containing 50mM NaH2PO4, 50mMNa2HPO4, 0.5mM K3Fe (CN)6, 0.5mM K4Fe(CN)6, 0.1% Triton X-100, 10mM Na2-EDTA, and 2 mM X-Gluc, pH 7.0, at 37°C. Samples were vacuum-infiltrated for 2 hours to 12 hours in staining solution, after which time they were washed with 75% ethanol until the tissues became clear. Images were collected under a stereomicroscope (KL300 LED, Leica) or light microscope (ICC50 HD, Leica).

### RT-PCR and qRT-PCR analysis

For expression analysis, total RNA was extracted using TRIzol solution (Invitrogen) and reverse-transcribed according to the manufacturer’s instructions (Invitrogen). The rice *ACTIN1* gene (Os03g50890) was amplified using the primers qActin-fw and qActin-rv as an internal standard to normalize the expression of studied genes such as *SHRs*, *IDDs* and *PINs*. qRT-PCR was performed using the iQ5 real-time PCR detection system (Bio-Rad) with real-time PCR Master Mix (TOYOBO, Japan). The primers used for expression analysis are described in Supplementary Table 4. The PCR conditions were as follows: pre-incubation at 94°C for 2 min followed by 40 cycles of 94°C for 20 sec, 62°C for 20 sec, and 72°C for 20 sec. All experiments were performed using three biological replicates (three different populations grown at different times) and three technical replicates.

### Leaf anatomy measurements

For quantification of leaf length, leaf width and leaf thickness, flag leaves of the first tiller of at least 11 independent plants were measured in the paddy field 30 days after flowering of Nipponbare or growth chamber 15 days after flowering of ME034V. For leaf anatomical traits, flag leaves of two tillers for each of the at least 11 independent lines were collected and hand sectioned. We recorded the number of mesophyll cells layers, numbers of mesophyll cells between veins, numbers of major veins, numbers of lateral and minor veins. To estimate bundle sheath, mesophyll cell and minor/intermediate vein size cross sections of a single leaf segment was taken from the center of the leaf blade, avoiding segments immediately adjacent to the midrib and lateral edges of the cross section. Quantification of cell, minor/intermediate vein area was performed with ImageJ software (NIH, https://imagej.nih.gov/ij/).

### *In situ* hybridization

*In situ* hybridizations were performed according to (Traas, 2008) with some modifications. Plant tissues were fixed in cold, freshly prepared FAA overnight at 4°C, dehydrated through an ethanol series (60%-75%-80%-95%-100%-100% with hourly transfers) at 4°C and kept in 100% ethanol for 2 d at -20°C. The samples were washed, infiltrated, sectioned, deparaffinized and rehydrated as described in (Traas, 2008). DIG-labelled sense and antisense probes to measure *OsSHR1*, *OsSHR2*, *OsIDD12*, *OsIDD13*, and *OsPIN5c* expression are shown in Supplementary Table 4. Because of the relatively weak expression of above genes, we performed a series of hybridization and washing treatments to optimize signal detection. The optimal hybridization temperature was performed at 40°C, which was much lower than previously reported. The slides were washed with 0.2X SSC two times at room temperature then 50°C at least 2 times. A BCIP/NBT kit (Roche, USA) was used for signal detection according manufacturer’s recommendations and expression visualized by photography under light microscopy.

### Laser capture microdissection (LCM)

The LCM method was used for isolation of SAM, P1 to P6 tissues from overexpression or knock-out transgenic lines using a modified protocol from (Takahashi et al., 2010) using a Veritas Laser Microdissection System LCC1704 (Molecular Devices, Leica). Selected areas were captured by an infrared laser (IR laser) onto CapSure Macro LCM Caps (Molecular Devices), and were subsequently cut by an IR laser. The target cells that fused to the LCM cap were collected by removing the cap from the tissue section and quickly frozen at -80°C prior to RNA isolations and reverse transcription.

### Yeast two-hybrid assay

The full-length coding region of *OsSHR1* was cloned into the *pGBKT7* vector as bait, and yeast two-hybrid screening was performed with a yeast expression library of rice cDNA (mRNA isolated from two-week-old rice seedling as described above, and library generated by Ouyi Biomedical Technology, Shanghai, China). Co-transformed yeast (*Saccharomyces cerevisiae*) AH109 (BD Biosciences, USA) strain according to the manufacturer’s protocol and plated on SD selective media lacking leucine, tryptophan, histidine and adenine (with 40 mM 3-AT). The positive clones were subsequently identified by sequencing. For confirmation of the specific interaction in yeast, the protein-coding regions of *OsSHR1*, *OsSHR2*, *OsIDD12* and *OsIDD13* were amplified using gene-specific primers (Supplementary Table 4), and the PCR products were fused into the activation domain (AD) vector *pGADT7* and/or the DNA-binding domain (BD) vector *pGBKT7* and verified by sequencing. The constructs were then co-transformed into yeast AH109 strain and plated on SD selective media and no detectable self-activation was observed (SD -His-Leu, + 2 mM 3-AT or SD -His-Trp, + 40 mM 3-AT (for OsSHR1-BD) or 20 mM 3-AT (for OsSHR2-BD)). Then, yeast cells were spotted on SD media lacking leucine, tryptophan, histidine and adenine, and their growth was compared after 2 to 4 days. Each assay was repeated three times.

### Protein-protein pull-down test

Full length cDNA of target protein OsSHR1, OsSHR2, OsIDD12 and OsIDD13 were amplified then ligated in *pGEX-6P-1* (GE Healthcare) and/or pLM303 (A derivative of *pET27a*, Novagen, USA) in frame fusion with GST and/or MBP respectively. GST-OsIDDs or MBP-OsSHRs fusion protein were expression in *E.coli* Rosetta strain, and fusion proteins purified according to manufacturer’s instructions (GE Healthcare or Novagen). Protein-protein binding buffer contained : 50 mM Tris-HCl (pH7.5), 100 mM NaCl, 0.25% Triton-X 100, 35 mM β-mercaptoethanol. Specific interactions were detected by western blotting with anti-MBP (Abcam) or anti-GST (Abcam) antibodies in proper dilution.

### BiFC assay

The protein-coding regions of *OsSHR1*, *OsSHR2*, *OsIDD12* and *OsIDD13* were amplified using gene-specific primers (Supplementary Table 4) and PCR products ligated into *pSAT1-nVenus-C* or *pSAT1-cCFP-N* to produce *pSAT1-nVenus-C-OsSHR1*, *pSAT1-nVenus-C-OsSHR2*, *pSAT1-cCFP-N-OsIDD12*, and *pSAT1-cCFP-N-OsIDD13*. All constructs were verified by sequencing. Plasmid pairs were transfected into rice protoplasts as described previously, using pairs of each construct and the empty vector as negatives control (Sun et al., 2016). The cells were then examined by laser scanning confocal microscopy (TCS SP2, Leica). YFP fluorescence was imaged by excitation with the 515-nm argon laser line and a 535– 565 nm band-pass emission filter.

### qPCR-based DNA pull-down/DNA binding assay

GST-OsIDD12, and GST-OsIDD13 recombinant proteins expression and purification was described above. DNA binding assay was performed based on previous description (Meng et al., 2013). The putative IDD proteins binding site presenting in *OsPIN5c* 1.4kb long intron 3 were designed fragments named F1 to F4 (Sequences presented in Supplementary), the amplified F1 to F4 fragments with equal amounts were incubated with GST alone or with the GST-OsIDDs fusion proteins, the DNA binding activity (protein bound DNA) was determined by qPCR after washing and elution. Primers used were listed in Supplementary Table 4.

### Electrophoretic mobility shift (EMSA) assay in vitro

The GST-OsIDD12 and GST-OsIDD13 recombinant protein expression and purification was performed as described above (Liu et al., 2008). The wild type IDD binding ‘TTTGTCGCTTT’ motif with its adjoining right 20 bp and left 20 bp sequence were synthesized and labeled with biotin as probe and different mutants of ‘TTTGTCGCTTT’ motif were also synthesized and labeled. The primers used are listed in Supplementary Table 4. The EMSA were performed as described (Liu et al., 2008) and visualized with ECL reagent (GE Healthcare).

### Chromatin immunoprecipitation (ChIP)-qPCR assay

The T_2_ generation of *pOsIDD12::GFP:OsIDD12* and *pOsIDD13::GFP:OsIDD13* OX transgenic seedlings at 20 DAG were used as material for ChIP-qPCR assay. Seedlings were harvested carefully including the shoot base and fully expanded fifth leaves were discarded. About two grams of plant material were harvested and were washed with ddH_2_O three times. Briefly, the assay was performed as described in Zhao et al. (Zhao et al., 2020) and anti-GFP antibodies (Abcam) were used to immuno-precipitate the DNA-IDDs complexes. qPCR primers (Supplementary Table 4) targeted the IDD binding motif (TTTGTCGCTTT) named P2 and two neighboring control DNA fragments named P1 and P3 in the third intron of *OsPIN5c* were tested. All experiments were performed using three biological replicates and three technical replicates. The ChIP-qPCR results were provided as relative enrichment of DNA fragments.

### Dual luciferase reporter assay

The CaMV *35S* promoter was inserted into the *pCambia*1300 (Hajdukiewicz et al., 1994), backbone using the *Eco*RI site, and then, 3XHA or 3XFLAG fragments were inserted into the vector using *Hind*III sites and the In-Fusion kit (Clontech) to generate basic and control effector vectors *p35S::HA* or *p35S::FLAG*, then we built a series effector vectors such as full-length CDS of *OsIDD12* or *OsSHR1*, and full-length CDS of *ODIDD13* or *OsSHR2* inserted into basic vector *p35S::HA* or *p35S::FLAG* respectively. An approximately 2.5 kb region upstream of *OsPIN5c* was cloned with or without insertion of intron3 (I3) sequences amplified and inserted into the pGREEN II 0800-LUC vector digested by *Hind*III and *Bam*HI, generating the reporter vectors *pI3-ProPIN5c::LUC* (Intro 3 ligated on the 5’ end of *PIN5c* promoter fragment) and *pProPIN5c::LUC*. Vector pGREEN II 0800-LUC carrying the *Renilla* luciferase (REN) gene driven by the CaMV 35S promoter was named pLUC in this study and used as an internal control reporter vector. Equal amounts of relative effector and reporter vectors were co-transformed into rice protoplasts and incubated in the dark at 28 °C for 16 h. The luciferase (LUC) activity was measured with the Dual-Luciferase Reporter Assay System Kit (Promega E1910) using the TriStar2 Multimode Reader LB942 (Berthold Technologies). Relevant primer sequences are listed in Supplementary Table 4.

Accession Numbers

*OsSHR1* (Os07g0586900/LOC_Os07g39820)

*OsSHR2* (Os03g0433200/LOC_Os03g31880)

*OsIDD12* (Os08g0467100/LOC_Os08g36390)

*OsIDD13* (Os09g0449400/LOC_Os09g27650)

*OsPIN5c* (Os08g0529000/LOC_Os08g41720)

## Acknowledgements

This research was supported by the Agricultural Science and Technology Innovation Program -G2P Program (2020YFE0202300) and partly supported by the National Key Research and Development Program of China (2020YFA0907603).

## Author contributions

XH. Sun, QM. Liu, SZ. Teng, C. Deng and HS. Li performed most experiments. ST. Wu and JX. Wu performed transformation of Setaria. YW. Wang contribute a lot of in situ assays. XA Cui, ZG. Zhang and W.P. Quick gave critical advice to experiments. T.P. Brutnell, XH. Sun and TG. Lu designed the experiments and wrote the manuscript.

## Competing interests

The authors declare no competing interests.

Materials & Correspondence. Indicate the author(s) to whom correspondence and material requests should be addressed.

XH. Sun will correspond to material requests.

## Tables

Tables information can be found at the Supplementary data file and Table IV text file.

## Figures

Figures information can be found at the picture file.

## Figure legends

Figures legends information can be found at the picture file.

## Statistical information

### Supplementary information

Supplementary information can be found at the Supplementary data file.

Reporting guidelines

